# Cyclo(Pro-Tyr) elicits conserved cellular damage in fungi by targeting the [H^+^]ATPase Pma1 in plasma membrane domains

**DOI:** 10.1101/2023.12.29.573620

**Authors:** D. Vela-Corcia, J. Hierrezuelo, A.I. Pérez-Lorente, P. Stincone, A.K. Pakkir Shah, A. Grélard, A. de Vicente, A. Pérez García, A. Loquet, D. Petras, D. Romero

## Abstract

Bioactive metabolites play a crucial role in shaping interactions among diverse organisms. In this study, we identified cyclo(Pro-Tyr), a metabolite produced by *Bacillus velezensis*, as a potent inhibitor of *Botrytis cinerea* and *Caenorhabditis elegans*, two potential cohabitant eukaryotic organisms. Based on our investigation, cyclo(Pro-Tyr) disrupts plasma membrane polarization, induces oxidative stress and increases membrane fluidity, which compromises fungal membrane integrity. These cytological and physiological changes induced by cyclo(Pro-Tyr) may be triggered by the destabilization of membrane microdomains containing the [H^+^]ATPase Pma1. In response to cyclo(Pro-Tyr) stress, fungal cells activate a transcriptomic and metabolomic response, which primarily involves lipid metabolism and Reactive Oxygen Species (ROS) detoxification, to mitigate membrane damage. This similar response occurs in the nematode *C. elegans*, suggesting that cyclo(Pro-Tyr) universally targets eukaryotic cellular membranes.

## Introduction

Specific inhibitory metabolites play a crucial role in shaping the dynamics of interkingdom interactions between soil bacteria and other organisms, such as fungi or nematodes^1–4^. In addition to affecting the abundance and composition of microbial populations in the soil, this chemical warfare between different kingdoms has far-reaching implications for the overall functioning and health of terrestrial ecosystems^5^. Moreover, the intricate interplay between bacteria, fungi, and nematodes mediated by these inhibitory metabolites underscores the complex and interconnected nature of soil ecosystems, highlighting the importance of understanding and harnessing these interactions for biotechnological applications.

Cyclic peptides are a relatively unexplored class of bioactive secondary metabolites that participate in interkingdom interactions within marine and soil ecosystems^6,7^. The smallest versions of these molecules are cyclic dipeptides that contain 2,5-diketopiperazine rings. Their structure provides stability and resistance to degradation, leading to long-lasting persistence in soil and extended periods of biological activity^8^. These compounds diversify their toxicity over other microorganisms, including fungi, nematodes or competing bacterial species, and may even induce apoptosis in human cancer cells^9–13^. Complex cyclic peptides have been suggested to induce significant disturbances within lipid bilayers, leading to the formation of disordered pores; in addition, some of these peptides disrupt bacterial communication systems^9^. Nonetheless, comprehensive knowledge of their precise mechanism of action and the host response remains elusive. These molecules are frequently synthesized by various organisms, including mammals, through the biosynthesis of amino acids; however, the available data indicate that approximately 90% of cyclic peptide producers are bacterial^14^. These molecules are typically specialized metabolites or byproducts generated by the terminal cleavage of peptides and primarily function as cell signaling molecules^15^.

In this study, we identified the cyclic dipeptide cyclo(Pro-Tyr) produced by *B. velezensis* CECT 8237 as a potent toxic molecule against different fungal strains. Using a sublethal dose that is not extensively destructive, we delineated the sequence of events that eventually damage the target cell and host response. Cyclo(Pro-Tyr) provokes a global oxidative burst responsible for the chemical deterioration of lipids or proteins, causing the membrane to lose functionality. Fungal cells mitigate this aggression through i) forming endosomes to eliminate the portion of the damaged membrane along with the dipeptide and ii) redirecting fungal lipid metabolism to help alleviate oxidative stress. The loss of fungal membrane functionality, along with the disorganization of membrane microdomains and the presence of the protein Pma1 in endosomes, led us to identify Pma1 as the potential main membrane target of the cyclodipeptide. We also observed this mode of action and cellular response in nematodes, suggesting that the activity is specifically universal toward eukaryotic membranes. The deterioration of membrane microdomains containing Pma1 is fascinating collateral insult that could extend the lifespan of previously abandoned fungicides.

## Results

### The cyclic dipeptide cyclo(L-Pro-L-Tyr) produced by *B. velezensis* exhibits potent antifungal activity

Butanol-extracted *B. velezensis* CECT 8237 cell-free supernatant was fractionated by silica gel column chromatography. A nonpolar mobile phase of chloroform:methanol (9:1) was used to minimize the presence of lipopeptides, and 72 fractions were collected. The activity of the fractions was assessed by measuring the level of oxidative stress (ROS) induced in germinated spores of *Botrytis cinerea* B05.10 (Fig. S1). Fraction 20 was the most active, provoking a ROS response significantly higher than that of hydrogen peroxide, which was used as a reference (P < 0.0001). Substances present in fraction 20 were further separated and purified via preparative HPLC with an elution gradient of water-acetonitrile. All the collected fractions (33 in total) were screened for antifungal activity to identify the putative bioactive compounds (Fig. S2). A higher ROS response was induced with fraction 24 (P < 0.0001) (Fig. 1a). Analysis of this fraction via LC-HRMS revealed the presence of three traces (Fig. 1b) with molecular weights of 185.12 Da 261.12 Da, and 521.23 Da, respectively (positive ion ESI-MS (m/z)), which corresponded to the double ion of molecule 2 (m/z of 261.12 Da). The two substances were retrieved in a database and compared with the standard spectrum. Detailed analysis of the gas phase fragmentation was performed by MALDI-ToF-MS/MS. Substance 1 was identified as a cyclodipeptide of L-Ala-L-Ile with a deduced molecular formula of C_9_H_16_N_2_O_2_. Substance 2 was identified as a cyclodipeptide of L-Pro-L-Tyr with a deduced molecular formula of C_14_H_16_N_2_O_3_ (Fig. 1c). The molecular formula was established on the basis of HR-ES-MS data ([M+H]^+^ at *m/z* 261,1254), and we confirmed the chemical structure of the cyclic peptide using solution nuclear magnetic resonance (NMR) spectroscopy (Fig. 1d). 1D proton (^1^H) and ^13^C experiments combined with 2D COSY, TOCSY and HMBC were used to identify ^1^H and ^13^C NMR resonances, confirming phenol resonances belonging to the Tyrosine residue (7.05 and 6.72 ppm) and the proline spin system.

**Figure 1.**
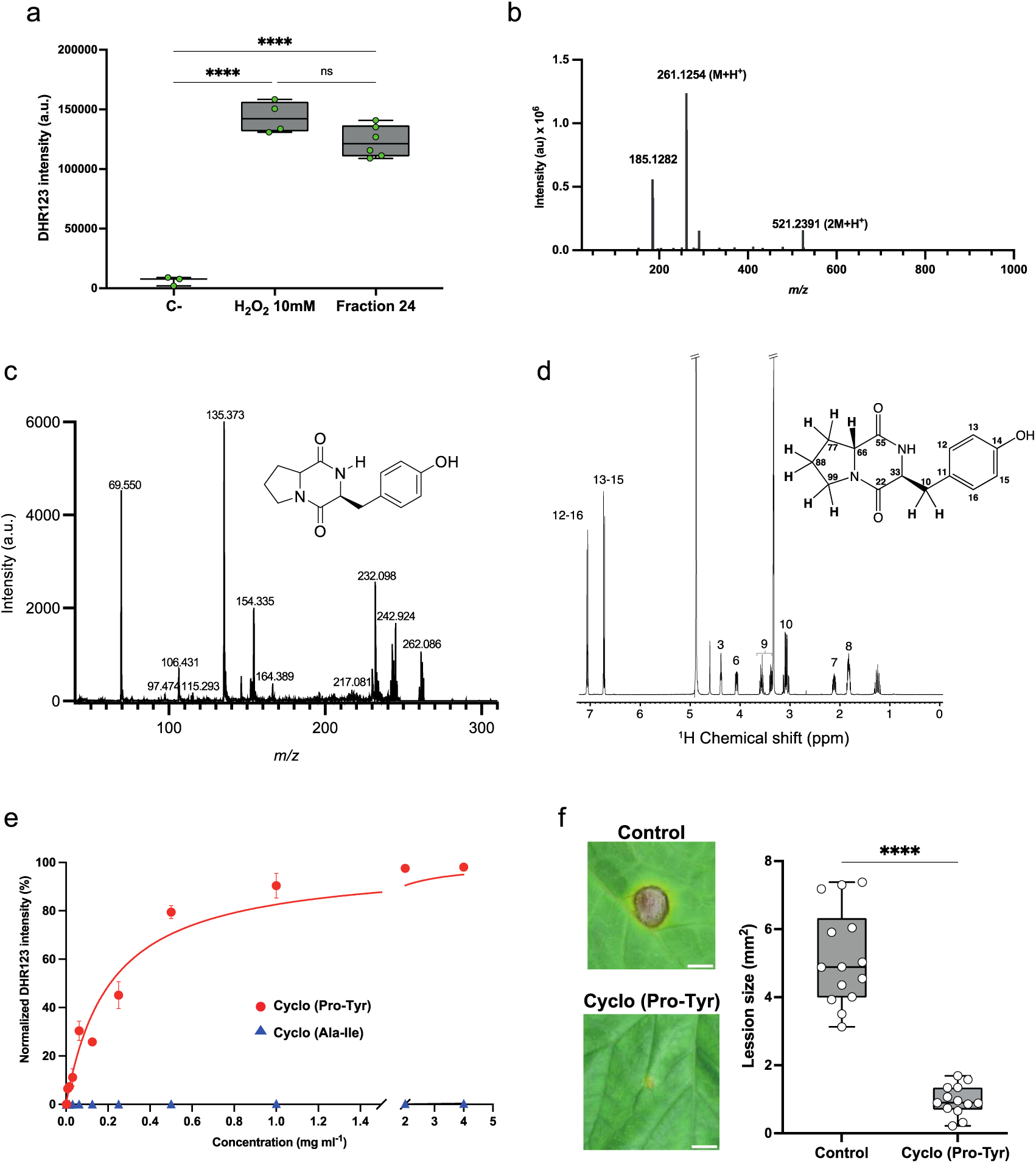
Isolation and identification of bioactive compounds present in *B. velezensis* CECT 8237 supernatant. **a**. The most active fraction obtained from supernatant fractionation was separated by HPLC and assayed against *B. cinerea* B05.10 germinated conidia. The highest activity was attained with peak 24, raising a significant ROS response (*P* < 0.0001) in comparison to the positive control with hydrogen peroxide. **b**. Peak 24 was analyzed by LC/HRMS, disclosing the presence of two different molecules, with masses of 185.12 Da and 261.12 Da, respectively, and a third peak corresponding to the double ion of molecule 2 with a m/z of 521.23 Da. **c**. Mass spectral fragmentation performed by MALDI-ToF-MS/MS identified substance 2 as a cyclodipeptide of L-Pro-L-Tyr with a deduced molecular formula of C_14_H_16_N_2_O_3_. The molecular formula was established by HR-ES-MS data ([M+H]^+^ at *m/z* 261,1254). **d**. Analysis by H^1^-NMR confirmed the structure corresponding to a cyclic dipeptide that contains aromatic signals from the phenol residue of a tyrosine lateral group (δ_H_ 7.04 and 6.81). A quartet of doublets (qd) at δ_H_ 3.06 can be assigned to the benzyl methylene protons of the Tyr residue, and a triplet at δ_H_ 4.35 corresponds to an adjacent CH. Finally, several multiplets of the aliphatic cycle at δ_H_ 1.23, 1.80, 2.09, 3.35 and 3.54 and a double doublet at δ_H_ 4.04 confirmed the heterocyclic structure. **e**. A concentration range of two different cyclodipeptides present in peak 24 was screened for ROS triggering as described above. Cyclo(Ala-Ile) showed no activity even at high concentrations, while Cyclo(Pro-Tyr) showed significant activity in a concentration-dependent manner. The half maximal effective concentration (EC_50_) was determined as 0.2 mg mL^-1^ for cyclo(Pro-Tyr). **f**. Lesion size triggered by *B. cinerea* on melon leaves after 72 hpi was reduced by 80% upon treatment with cyclo(Pro-Tyr). Whiskers’ plot shows all measurements (green dots), medians (black line), and minimum and maximum (whiskers ends). Data sets did not pass the Shapiro‒Wilk test for normality (P > 0.05) and were compared using a nonparametric two-tailed Mann‒Whitney test, with quadruple asterisks indicating significant differences at P < 0.0001.

Different concentrations of purified cyclodipeptides were screened for antimicrobial activity through ROS triggering on *B. cinerea* spore suspension. Only the cyclodipeptide (Pro-Tyr) induced a significant ROS response to the fungal spores in a concentration-dependent manner. The EC_50_ that could induce an oxidative burst in 50% of the total response was 0.25 mg mL^-1^ cyclo(Pro-Tyr) (Fig. 1e). A bioassay conducted by inoculating the third leaf of melon plants with a spore suspension of *B. cinerea* confirmed this *ex vivo* antifungal activity. After the third leaf of melon plants was sprayed with a 0.25 mg mL^-1^ cyclo(Pro-Tyr) solution, the leaves were inoculated with *B. cinerea,* and lesions were recorded 72 h post-inoculation. Treatment with cyclo(Pro-Tyr) reduced the necrotic lesions induced by the fungus by 81.2% in comparison with untreated plants (Fig. 1f).

### Foci of cyclo(Pro-Tyr) in the fungal membrane correlate with the formation of endosomes and membrane depolarization

The induction of ROS led us to study the ultrastructural changes in *B. cinere*a hyphae via transmission electron microscopy. The most noticeable cytological alterations were concentrated in the cell membrane and predominantly involved the formation of small membrane-bound compartments called endosomes, which are associated with peroxisomes (Fig. 2a). Early endosomes have been reported to mediate the internalization of plasma membrane sections by endocytosis^16^, leading to a loss in membrane integrity and increasing the vulnerability of hyphae to environmental stresses^17^. To further clarify the subcellular location of cyclo(Pro-Tyr), the molecule was synthesized and conjugated to fluorescein by direct synthesis, and this conjugated derivative was traced on fungal cells via confocal laser scanning microscopy (CLSM). According to the cytological alterations, green fluorescent foci (cyclopeptide-FL) were regularly distributed along the cell membrane (Fig. 2b, arrowhead). Double staining with the endocytic marker dye FM 4-64, which is taken into the cell by endocytosis and accumulates in the vacuolar tonoplast, revealed that cyclodipeptide foci were colocalized with the inner part of the plasma membrane. These findings support that the cyclodipeptide was internalized into the cells via endocytosis and subsequently transported into early endosomes (Fig. 2b). Through using a Kymograph, we obtained information on the movement of early endosomes containing cyclic dipeptide, which moved in a retrograde direction toward older sections of the hypha (Fig. 2c and Video S1). In contrast to the hyphae treated with the cycle-dipeptide, no vesicle movement was observed in untreated fungal hyphae (Video S2).

**Figure 2.**
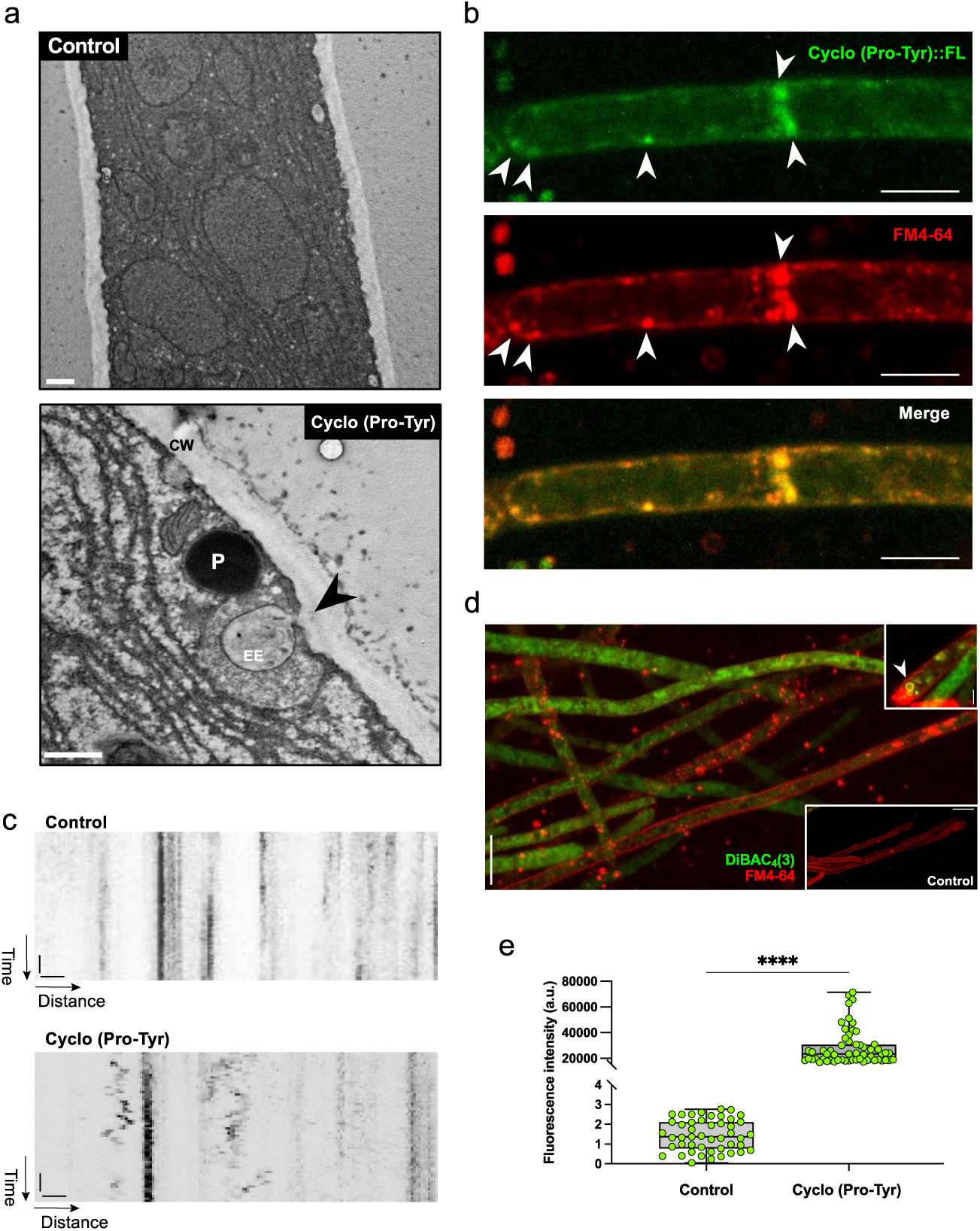
Anatomical damage elicited by cyclo(Pro-Tyr) and subcellular localization. **a**. Electron micrographs of the plasma membrane of untreated and cyclodipeptide-treated *B. cinerea*. Cyclo(Pro-Tyr) caused the formation of endosomes that internalize plasma membrane sections. These endosomes were attached to the membrane and distributed along the cyclodipeptide-treated hyphae; they were accompanied by a peroxisome to degrade the endosome content. Scale bar 0.5 μm. **b**. Fluorescent cyclo(Pro-Tyr) is localized, forming accumulations in foci at the inner part of the *B. cinerea* plasma membrane, and colocalized with endocytic vesicles counterstained with FM4-64. Scale bar 5 μm. **c**. Kymographs showing the dynamic behavior of endocytic vesicles in *B. cinerea* treated with the solvent DMSO (Control) and with 250 μg ml^−1^ cyclo(Pro-Tyr). Intracellular vesicle trafficking increased in the presence of cyclo(Pro-Tyr). Horizontal bar: 1 μm, vertical bar: 1 s. **d**. Cyclodipeptide-treated *B. cinerea* cells costained with DiBAC_4_(3) (green) and FM4-64 (red). Several cells are depolarized, as indicated by the uptake of the voltage-sensitive dye DiBAC_4_(3). Less green fluorescence was observed in untreated cells, denoting less depolarized cells (bottom inset). Likewise, endocytic vesicles formed after cyclo(Pro-Tyr) was added are also stained green, suggesting that the fungus internalizes depolarized membrane sections (top inset). Scale bar represents 5 μm. **e**. Whiskers plot showing the fluorescence intensity of *B. cinerea* cells stained with DiBAC_4_(3) after treatment with cyclo(Pro-Tyr). Whiskers’ plot shows all measurements (green dots), medians (black line), and minimum and maximum (whiskers ends). Data sets did not pass the Shapiro‒Wilk test for normality (P > 0.05) and were compared using a nonparametric two-tailed Mann‒Whitney test, with quadruple asterisks indicating significant differences at P < 0.0001. CW: Cell wall, P: Peroxisome, EE: Early endosome.

Based on the specific focal localization of the cyclopeptide and the fungal cellular response, the destabilization of certain membrane domains is the most likely mode of action. Thus, we evaluated the integrity of the plasma membrane by monitoring ion permeability in cyclodipeptide-treated cells. Eukaryotic cells selectively pump ions across their plasma membrane, which generates an electric potential^18^; thus, depolarization of the cell caused by an increase in ion permeability can be monitored by cellular accumulation of the anionic voltage-sensitive green fluorescent oxonol DiBAC_4_(3)^18,19^. We found that cyclo(Pro-Tyr) increased the number of DiBAC_4_(3)-positive *B. cinerea* cells compared to untreated cells (Fig. 2d, 2e), a finding indicative of membrane depolarization. Interestingly, endosome formation was concentrated on depolarized plasma membrane sections upon cyclodipeptide treatment (Fig. 2d, upper inset).

### Cyclo(Pro-Tyr) induces oxidative stress that impairs fungal membrane functionality by malfunctioning [H^+^]ATPase Pma1

Upon staining with the ROS fluorescent reporter dihydrorhodamine-123 (DHR-123), fluorescent signal emissions were specifically localized in mitochondria. This result indicated that the hyphae of *B. cinerea* showed high levels of oxidative stress, consistent with membrane dysfunctionality caused by the cyclopeptide (Fig 3a, b). These findings contrasted with the low intensity of the DHR-123-related signal emitted by untreated *B. cinerea* hyphae (Fig. 3a, inset). ROS target different cellular components, including membrane lipids^20,21^; thus, we analyzed the redox state of the plasma membrane by shifting the fluorescence emission of BODIPY™ 581/591 C11. An increase in the green fluorescence signal was observed in hyphae treated with cyclo(Pro-Tyr), indicating that the plasma membrane lipids undergo significant peroxidation (Fig. 3c-d). A consequence of lipid peroxidation is the increase in membrane fluidity^21,22^, which can be evaluated by monitoring the cellular distribution of red fluorescent lipophilic DilC_12_(3). Aggregates of this dye tend to move laterally along the plasma membrane, and changes in membrane fluidity result in differential accumulation^23^. According to a more fluid membrane, the treatment with the cyclodipeptide induced the formation of dye-aggregate foci, and untreated cells, which should present a less fluidified membrane, were homogeneously decorated with the fluorescent probe (Fig. 3e-f).

**Figure 3.**
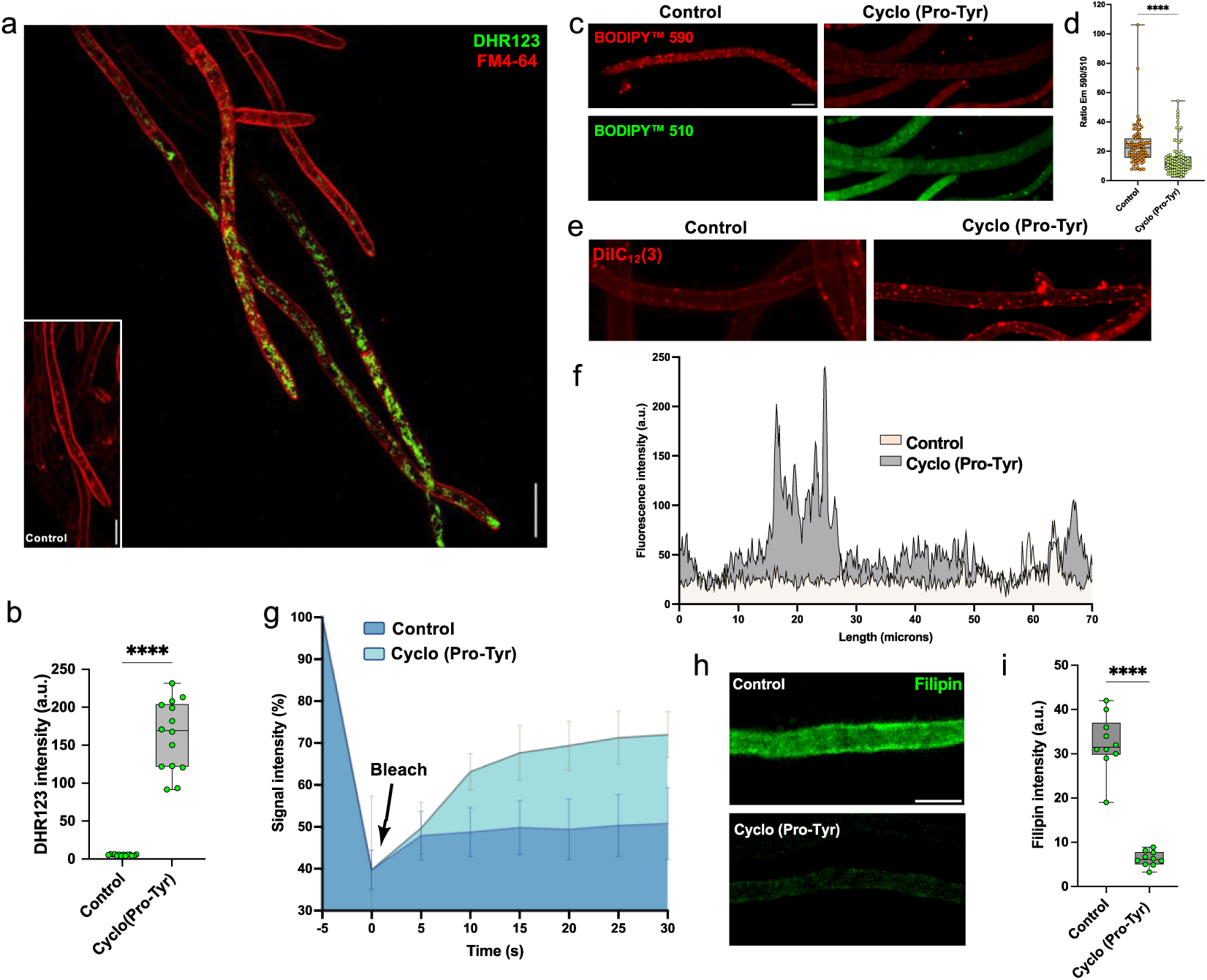
Effect of cyclo(Pro-Tyr) on the *B. cinerea* B05.10 plasma membrane. **a**. Oxidative burst triggered by cyclo(Pro-Tyr) in *B. cinerea* attained by using the ROS indicator dihydrorhodamine-123 (DHR-123). Upon addition of cyclodipeptide, mitochondria exhibited a very intense green fluorescence signal due to the oxidation of DHR123 to R123 elicited by ROS. **b**. Whisker plot showing the intensity of *B. cinerea* mitochondria stained with DHR123 after treatment with cyclo(Pro-Tyr). All measurements (green dots), medians (black line), and minimum and maximum (whisker ends) are represented. Data sets did not pass the Shapiro‒Wilk test for normality (P > 0.05) and were compared using a nonparametric two-tailed Mann‒Whitney test, with quadruple asterisks indicating significant differences at P < 0.0001. **c**. Representative fluorescence microscopy images of *B. cinerea* cells undergoing lipid peroxidation upon the addition of cyclo(Pro-Tyr). BODIPY-510 (oxidized) represents the levels of lipid ROS, while BODIPY-590 represents the level of staining with the probe (unoxidized). **d**. Whisker plot showing the 590/510 fluorescence emission ratio of *B. cinerea* BODIPY-stained cells after treatment with cyclo(Pro-Tyr). All measurements (orange and yellow dots), medians (black line), and minimum and maximum (whiskers ends) are represented. Data sets did not pass the Shapiro‒Wilk test for normality (P > 0.05) and were compared using a nonparametric two-tailed Mann‒Whitney test, with quadruple asterisks indicating significant differences at P < 0.0001. **e**. Representative fluorescence microscopy images showing the lateral distribution of red fluorescent lipophilic dye. Untreated *B. cinerea* cells show a homogeneous distribution along the hypha, whereas in cyclodipeptide-treated cells, the distribution tends to form large accumulations because the mobility along the plasma membrane was larger. **f**. Representative fluorescence intensity of DilC_12_(3) aggregates over *B. cinerea* hyphae. Upon treatment with cyclo(Pro-Tyr), the dye tends to form large aggregates due to the increased membrane fluidity. Untreated cells show a more homogeneous dye distribution along the hypha with less intense dye accumulation points **g**. Charts showing fluorescent recovery in a photobleached region of the plasma membrane of *B. cinerea*. Data points represent the mean ± standard error of the mean (SEM). After the bleaching event, cyclodipeptide-treated cells recovered up to 70% of the initial fluorescence because their membrane was more fluent, while control cells reached 45% of the initial fluorescence. **h**. Changes in the fluorescence pattern of filipin staining in *B. cinerea* hyphae under the effect of cyclo(Pro-Tyr) in comparison to untreated cells. Scale bar = 5 μm. **i**. Whisker plot showing the intensity of filipin fluorescence after treatment with cyclo(Pro-Tyr). All measurements (green dots), medians (black line), and minimum and maximum (whisker ends) are represented. Data sets passed the Shapiro‒Wilk test for normality (P > 0.05) and were compared using a parametric two-tailed Student’s *t* test with Welch’s correction, with quadruple asterisks indicating significant differences at P < 0.0001.

To further confirm this finding, we monitored the lateral mobility of membrane components via fluorescence recovery after photobleaching (FRAP) experiments. The recovery of the FM4-64 fluorescence signal occurred progressively in untreated and cyclodipeptide-treated cells. However, compared to untreated cells, cells treated with cyclodipeptides recovered the fluorescence signal more quickly and extensively (Fig. 3g), which is consistent with the increased lateral movement of membrane components due to the increased plasma membrane fluidity. The fluidity of the plasma membrane is regulated by diverse factors, including the lipid composition of the membrane, the presence of membrane proteins, and the level of sterols^24–27^ that regulate the packing and ordering of lipid molecules^28,29^. Thus, we examined the level of accumulation of ergosterol in the fungal plasma membrane using the specific fluorescence dye filipin. Filipin emits a green fluorescence signal upon binding to ergosterol, and a correlation can be established between the intensity of filipin fluorescence and the level of ergosterol in the plasma membrane. Compared to untreated cells, treated cells showed a comparatively low level of fluorescent signal, which might result from a reduction in sterol levels or the diffusion of ergosterol along the fungal plasma membrane in response to cyclodipeptide activity (Fig. 3i); these cellular responses are different but not exclusive.

Inspired by these findings and the depolarization of the cell membrane, we used immunocytochemistry to investigate the localization and abundance of Pma1, an encoding plasma membrane [H^+^]ATPase. No detectable signal was observed in untreated *Botrytis* hyphae, which might indicate a protein membrane content below the detectable threshold (Fig. 4a). Corresponding with endosome formation in response to cyclodipeptide (Fig. 4a), discrete fluorescent foci related to Pma1 colocalized with FM4-64-labeled endosomes in treated fungal cells. Similar findings were obtained with the fungal pathogen *Candida albicans* that expressed the translational fusion Pma1-GFP (Fig. 4b). Molecular docking analysis resulted in 36 clusters of putative binding sites distributed across six different regions of the Pma1 protein. Accordingly, the most energetically favorable binding site was located within the C-terminal domain of Pma1, precisely within the cavity formed by alpha-helices M5, M7, M8 and M10 (Fig. 4c). The binding of cyclo(Pro-Tyr) to Pma1 relies on the establishment of four hydrogen bonds with the residues K749, A814, Y875, and Q879. These interactions featured remarkably close distances, measuring 2.44 Å, 1.88 Å, 2.13 Å, and 2.25 Å, respectively (Fig. 4d).

**Figure 4.**
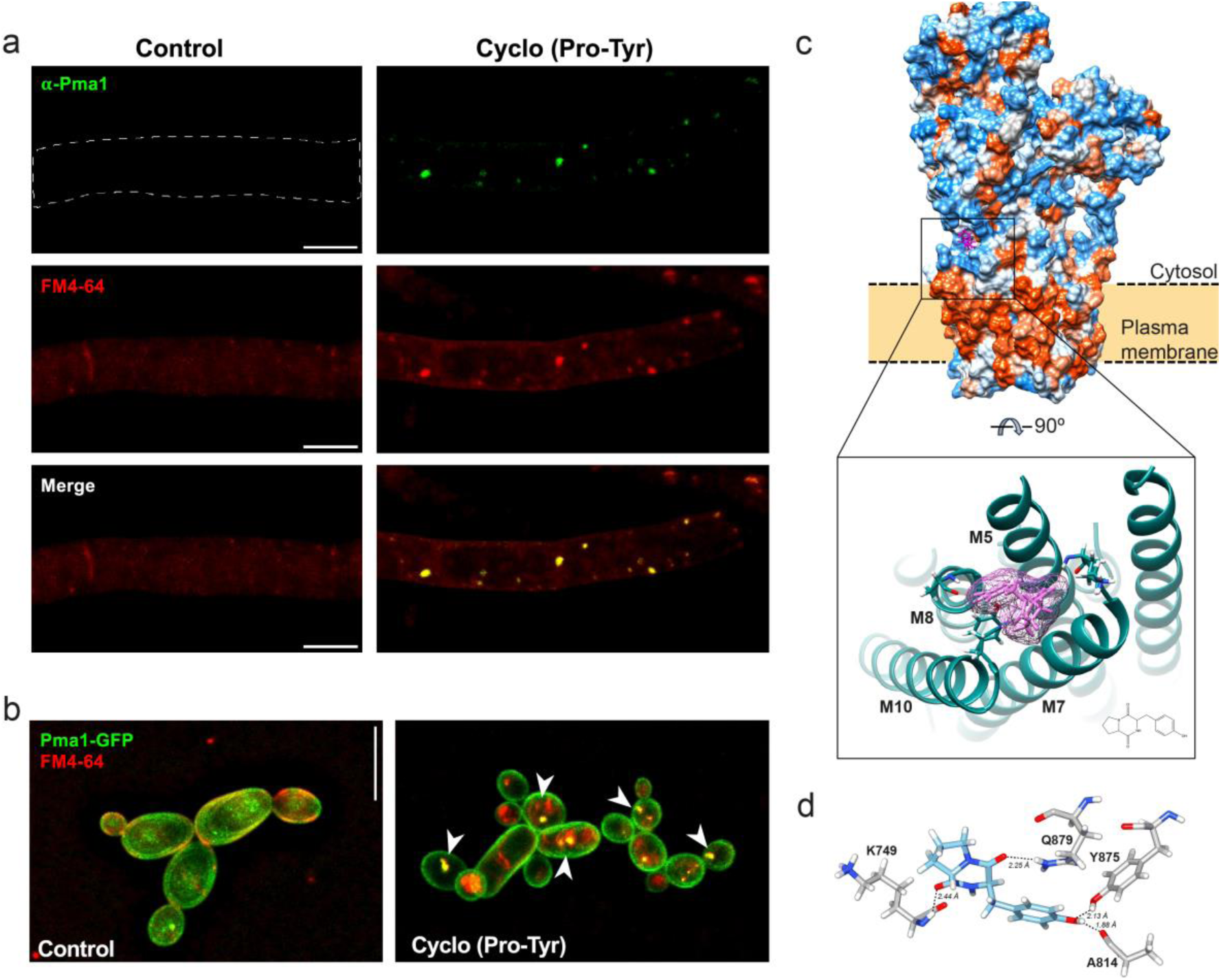
Effect of cyclo(Pro-Tyr) on membrane microdomains. **a**. Representative fluorescence microscopy images of Pma1 immunodetection in *B. cinerea* cells treated with cyclo(Pro-Tyr) in comparison with untreated cells. Immunocytochemistry using an anti-Pma1 mouse monoclonal primary antibody and an anti-mouse GFP-conjugated secondary antibody showed that Pma1 formed foci in *B. cinerea* cells that colocalized with endocytic vesicles stained with FM4-64 upon the addition of cyclo(Pro-Tyr). No Pma1 signal was observed in control cells. Scale bar 5 μm. **b**. Representative fluorescence microscopy images of GFP-tagged Pma1 localization in *C. albicans*. In control cells, Pma1 localization was restricted to the plasma membrane, and no internal vesicles stained with FM4-64 were observed. In cyclodipeptide-treated cells, Pma1 localized in the plasma membrane as well as intracellular foci that colocalized with endocytic vesicles (arrowheads). Scale bar, 10 μm. **c**. Structural model of the Pma1 monomer subunit depicted as hydrophobic potential showing the insertion within the plasma membrane and the putative interaction with cyclo(Pro-Tyr). The putative binding site was formed by the cavity between alpha helices M5, M7, M8 and M10 located at the C-terminal domain. **d**. Molecular docking between the cyclo(Pro-Tyr) molecule and Pma1. The proposed binding site is formed by the residues K749, A814, Y875, and Q879. The measured distances between the cyclo(Pro-Tyr) and side chains of these residues were 2.44 Å, 1.88 Å, 2.13 Å, and 2.25 Å, respectively.

To rule out that the effect of cyclo(Pro-Tyr) is a result of a direct lipid membrane destabilization, we investigated the impact of cyclodipeptide addition on reconstituted lipid vesicles. Multi-lamellar vesicles were reconstituted using a lipid composition mimicking fungal membranes (PC/PE molar ratio of 70/30, including deuterated POPC-d_31_) and probed by solid-state NMR (Fig. S3). Static ^2^H NMR experiment of PC/PE vesicles reveals a characteristic spectral pattern made of well-resolved Pake doublets, typical for a lamellar phase. In presence of cyclo(Pro-Tyr) at a peptide-to-lipid molar ratio of 1/20, no spectral broadening was detected, suggesting that the peptide has no significant effect of the lipid membrane fluidity. Extraction of the structural order parameters for each carbon position along the acyl chain of the lipids revealed no perturbation induced by the cyclopeptide addition. In addition, ^13^C-detected magic-angle spinning solid-state NMR experiments were used to probe the mobility of the dicyclopeptide in the membrane. Peptide signals were not detected in the cross-polarization spectrum and unambiguously detected in the INEPT spectrum, suggesting that the peptide is very weakly bound to the membrane. Altogether, these results suggest that it is not the cyclopeptide per se that directly destabilizes and damages the lipid membrane.

After cyclo(Pro-Tyr) was added, a significant increase in the number of endosomes was measured in *Candida albicans*-treated cells compared to untreated cells (Fig. S4a). This observation suggests that the effect of the cyclodipeptide on the formation of endosomes containing Pma1 in non-phylogenetically related fungal species was conserved. Pma1 is localized in the membrane domains called lipid rafts; thus, we hypothesized that disorganization of this microdomain results from the relief of Pma1. DRM-associated proteins were isolated from cyclo(Pro-Tyr)-treated or untreated cells, resolved in SDS gels and revealed with anti-Pma1 primary antibody at 12 h and 24 h after treatment. An immunoreactive band for Pma1 was clearly detected in the DRM fraction of untreated cells, with a maximum intensity at 24 h. However, treatment with cyclo(Pro-Tyr) provoked a reduction in signal intensity, indicative of the lower quantity of Pma1 (Fig. S4b); this reduction mostly likely resulted from the internalization events of membrane domains promoted by the cyclodipeptide.

### The lipid metabolic pathway is mainly adjusted in *B. cinerea* in response to cyclo(Pro-Tyr) toxicity

Cyclo(Pro-Tyr) modifies the dynamics and functionality of the fungal cell membrane, which, along with all cellular modifications, led us to explore rearrangements at the genetic or metabolic level. Twenty-four hours after cyclo(Pro-Tyr) was added, the whole transcriptional analysis of fungal hyphae revealed that the expression of 295 genes was induced and 246 genes were repressed (Fig. 5a). Unfortunately, most of these differentially expressed genes were annotated as uncharacterized proteins, challenging further analysis and interpretation of the results. The annotated induced genes were involved in very specific biological processes, including carbohydrate metabolism, secondary metabolite biosynthesis or stress response. The classification of the induced genes in the Kyoto Encyclopedia of Genes and Genomes (KEGG) pathway and Gene Ontology (GO) terms showed that pathways related to the biosynthesis of fatty acids and glycosphingolipids were specifically overrepresented (Fig. 5b), which might be related to the dynamics of the fungal cell membrane. Phosphatidylserine decarboxylase (*bcpsd*) is dedicated to the biosynthesis of phosphatidylethanolamine, which is an elemental component of cellular membranes. Our cytological study showed that the level of ergosterol in the cell membrane was low; however, we found that *erg28* was overexpressed, which is part of the ergosterol biosynthesis pathway. The detoxification of xenobiotics was activated, leading to the induction of an acyl-coenzyme A synthetase specific for medium-chain fatty acids. These enzymes catalyze fatty acid activation, the first step of fatty acid metabolism, through transferring acyl-CoA and participate in the glycine conjugation pathway during the detoxification of xenobiotics. In addition, we found mono- and dioxygenases, which are part of cellular detoxification programs. Finally, the upregulation of a hydroxysteroid dehydrogenase, which coincided with the association between peroxisomes and endosomes (Fig. 2a), suggested that the breakdown of fatty acids was activated by peroxisomal β-oxidation.

**Figure 5.**
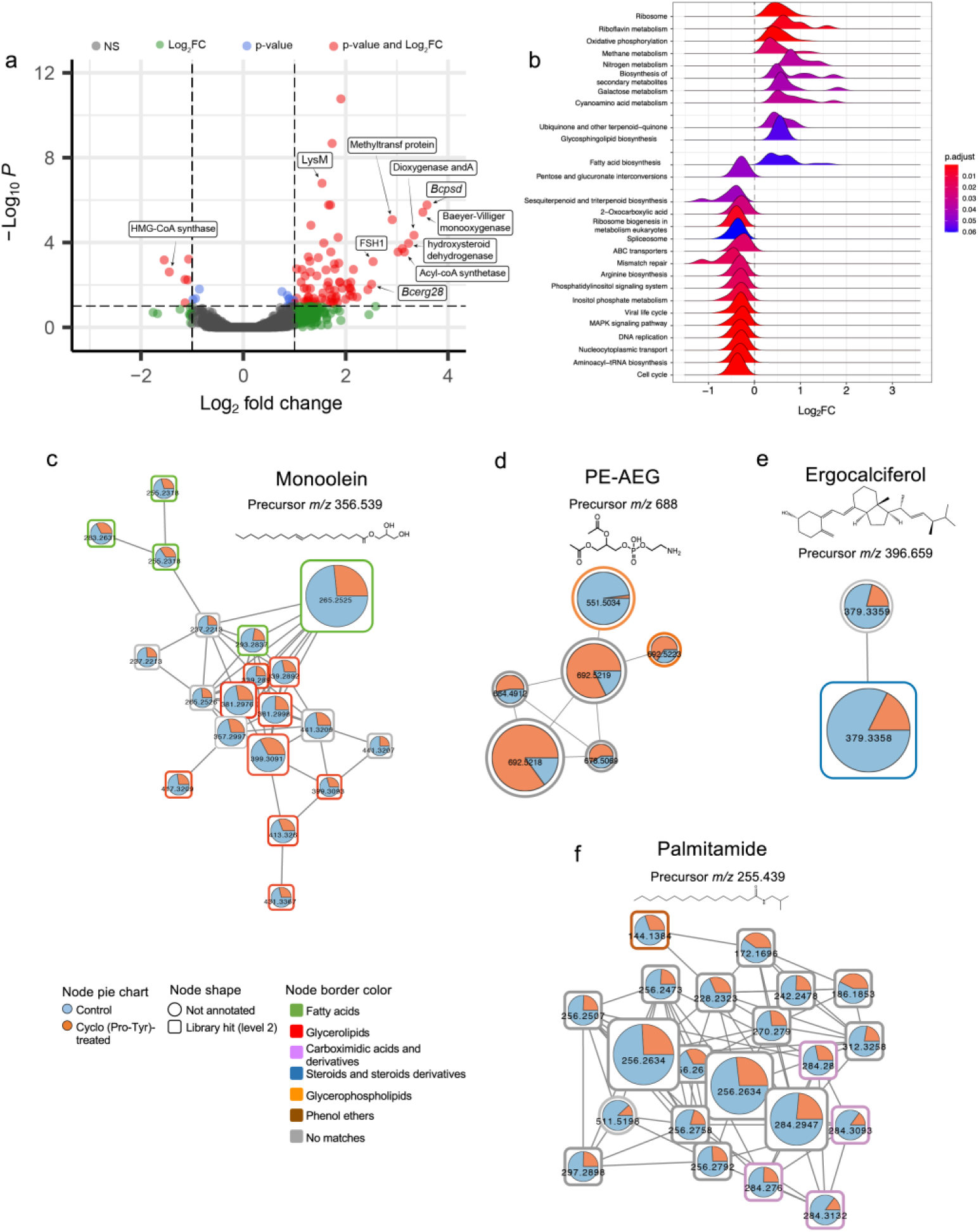
Response of *B. cinerea* B05.10 following treatment with cyclo(Pro-Tyr). **a**. Volcano plot of DEGs identified by total transcriptome analysis of cyclodipeptide-treated and untreated cells 24 h after treatment. P values were calculated on the basis of the Fisher method using nominal P values provided by edgeR and DEseq2. Dashed lines represent the threshold defined for P (horizontal) and fold change (vertical) for a gene to be considered a DEG. Tags label the genes related to plasma membrane composition and xenobiotic compound detoxification: *Bcpsd*, phosphatidylserine decarboxylase; mono- and dioxygenase, detoxification of xenobiotic compounds; Acyl-CoA synthetase, catalyzes fatty acid activation; hydroxysteroid dehydrogenase, peroxisomal β-oxidation of fatty acids. NS, not significant. **b**. Transcriptional analysis of DEG KEGG pathways in *B. cinerea* after treatment with cyclo(Pro-Tyr) for 24 h. Metabolic pathways related to fatty acid biosynthesis and glycosphingolipid biosynthesis were induced upon challenge with cyclo(Pro-Tyr). **c**. Molecular families corresponding to monoolein; phosphatidylethanolamine conjugated to acyl ether glycerol (PE-AEG); ergocalciferol; or palmitamide. Pie charts represent the mean peak abundance of metabolites in each condition. Node shape indicates the level of identification according to reference. Node border color indicates the chemical class of each metabolite.

To further confirm these observations, we studied the metabolic profiles of fungal cells after treatment with cyclo(Pro-Tyr). In agreement with our transcriptome analysis, partial least-squares discriminant analysis (PLS-DA) revealed that the discriminating metabolites differentially accumulated in hyphae treated with cyclo(Pro-Tyr) were fatty acyls, organooxygen compounds or carboxylic acids (Fig. S3). The complementary use of SIRIUS/ZODIAC, CSI:FingerID and CANOPUS predicted molecular formulas, compound class and putative chemical structure based on *in silico* MS/MS fragmentation trees. The results of principal component analysis (PCA) and heatmap analyses indicated that the samples were clustered in two groups (Fig. S5a-b), and (PLS-DA) identified fatty acyls and carboxylic acids as the differentially accumulated metabolites after treatment with cyclo(Pro-Tyr) (Fig. S5c). Through comparing cyclodipeptide-treated and untreated cells, we discovered a significant depletion of monoolein (*m/z* 356.539), a polyunsaturated phospholipid critical for maintaining the structural integrity and fluidity of the plasma membrane. In contrast, we observed an increase in saturated phosphatidylethanolamine conjugated with acyl and ether (PE-AEG) (*m/z* 688.895) (Fig. 5c,d). Phosphatidylethanolamine is involved in the synthesis and maintenance of lipid components in the cell membrane; therefore, changes in the accumulation levels of these phospholipids will modify the function and permeability of the plasma membrane, which ultimately determine the wealth and survival of the fungal cell.

Consistent with the lower ergosterol content in the membranes of hyphae treated with cyclo(Pro-Tyr), our metabolic analysis showed that ergocalciferol was depleted (*m/z* 396.659). In fungi, ergosterol serves as a precursor to ergocalciferol, which is further converted to the active form of vitamin D, a molecule vital for fungal growth and development. In addition, ergosterol and its derivatives, including ergocalciferol, are important components of fungal cell membranes. Thus, depletion of ergocalciferol might modify the structure and function of the cell membrane, influencing the overall fungal biology and physiology (Fig. 5e). Finally, a noticeable decrease in the levels of palmitamide was observed in treated cells. Palmitamide is a lipidic molecule with remarkable antioxidant and anti-inflammatory properties. It effectively shields cells from cytotoxicity by significantly reducing the production of ROS, an action that helps maintain cellular homeostasis under oxidative stress. The modulation of palmitamide levels in response to cyclo(Pro-Tyr) may indicate that palmitamide is involved in the intricate network of lipid signaling and highlights the potential therapeutic implications of this lipid compound in managing oxidative disorders (Fig. 5f).

### Cyclo(Pro-Tyr) reproduced cytological and physiological symptoms in nematodes

Based on the mode of action of cyclo(Pro-Tyr) at the cellular membrane, we hypothesized that other eukaryotes are potential targets. *Bacillus* coexists in soil with nematodes, and some are important parasites of plants. We had previously demonstrated that *Bacillus* cell-free supernatant protected three-week-old adult melon plants against infection by the root-knot nematode *Meloidogyne incognita* (Fig. S6a). In vitro experiments against *Caenorhabditis elegans* strain Bristol N2 demonstrated that the nematocidal activity of the supernatant was independent of the main lipopeptides produced by *Bacillus* cells (Fig. S6b), suggesting the presence of alternative molecules responsible for this biological activity. The same fractions tested against *Botrytis* revealed that fraction 20, which contained ^30–33^cyclo(Pro-Tyr), was the most active fraction and could trigger a significant accumulation of ROS in *C. elegans* compared to the untreated fraction (Fig. S7a,b). We further aimed to determine the main nematode tissue targeted using cell lines fluorescently labeled in different parts of the digestive apparatus, from the pharynx to the intestine, and in intestinal epithelial cell junctions (Fig. S7c-e; Table S1). Compared to the homogenous signal distribution in the whole intestinal tract of untreated nematodes, observed in the line CL2122, fluorescence emission in this line was significantly impaired following treatment with cyclodipeptide (Fig. 6a-b). To further confirm the specific toxicity to the intestine, a study using propidium iodide was performed to establish a correlation between the decay of green fluorescence signal (representative of tissue damage) and an increase in red fluorescence emitted by propidium iodide (representative of cell death). However, no colocalization of either signal was observed.

**Figure 6.**
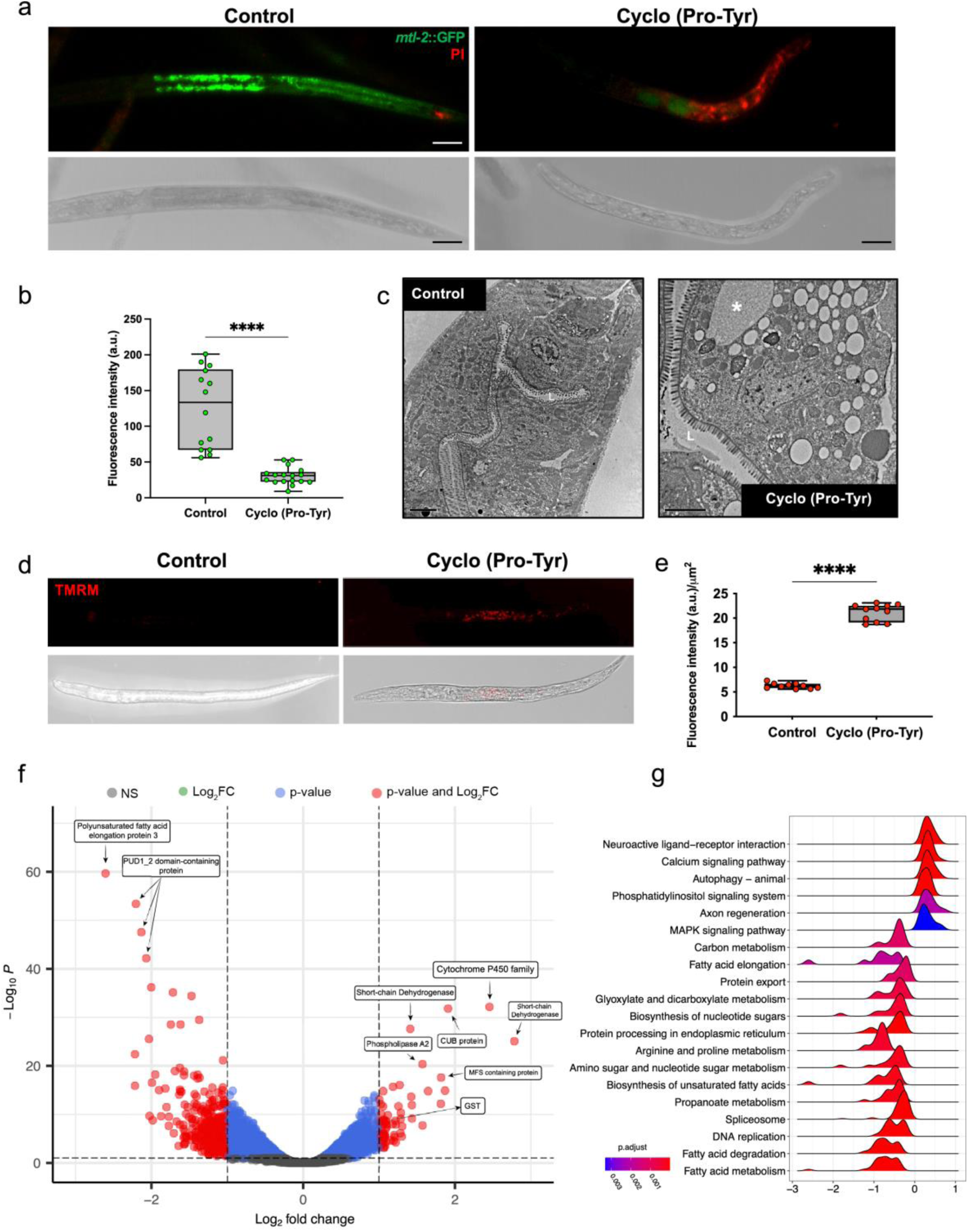
Response of *C. elegans* following treatment with cyclo(Pro-Tyr). **a**. Representative fluorescence microscopy images of *C. elegans* line CL2122, labeled in the intestinal epithelium cells, co-stained with propidium iodide (indicative of cell death). Scale bar 20 μm. **b**. Whisker plot showing the intensity of GFP fluorescence after treatment with cyclo(Pro-Tyr). All measurements (green dots), medians (black line), and minimum and maximum (whisker ends) are represented. Data sets did not pass the Shapiro‒Wilk test for normality (P > 0.05) and were compared using a nonparametric two-tailed Mann‒Whitney test, with quadruple asterisks indicating significant differences at P < 0.0001. **c**. Electron micrographs for the intestinal tract of untreated or cyclodipeptide-treated *C. elegans*. Cyclo(Pro-Tyr) caused an increase in vacuolization (asterisk) and endosome formation associated with peroxisomes. Scale bar, 2 μm. **d**. Representative fluorescence microscopy images of mitochondrial membrane potential, visualized with TMRM, in *C. elegans* N2 cells treated with cyclo(Pro-Tyr). Scale bar = 10 μm. **e**. Bar chart showing the effect of cyclo(Pro-Tyr) on TMRM signal intensity, which reflects the mitochondrial proton gradient in *C. elegans* N2 cells. Data sets did not pass the Shapiro‒Wilk test for normality (P > 0.05) and were compared using a nonparametric two-tailed Mann‒Whitney test, with quadruple asterisks indicating significant differences at P < 0.0001. **f**. Volcano plot of DEGs identified by total transcriptome analysis of cyclodipeptide-treated and untreated cells 24 h after treatment. P values were calculated on the basis of the Fisher method using nominal P values provided by edgeR and DEseq2. Dashed lines represent the threshold defined for P (horizontal) and fold change (vertical) for a gene to be considered a DEG. Tags label the genes related to plasma membrane composition and xenobiotic compound detoxification: Cytochrome P450, involved in the cholesterol biosynthesis pathway; short-chain dehydrogenases, involved in the aerobic biodegradation process; CUB-like domain containing protein, involved in tissue repair and toxic sensing; phospholipase A2 (PLA2), responsible for free polyunsaturated fatty acid (PUFA) generation; and glutathione S-transferase (GST), which catalyzes the conjugation of reduced glutathione. **g**. Transcriptional analysis of DEG KEGG pathways in *C. elegans* N2 after treatment with cyclo(Pro-Tyr) for 24 h. Pathways related to lipid metabolism were repressed. In contrast, signal transduction pathways were induced. L: Intestinal Lumen.

As previously described for *B. cinerea*, an anatomical study of nematodes treated with cyclo(Pro-Tyr) did not reveal drastic changes in the outer layer structures of the nematode tissue; the microvilli of the epithelial cells remained unaffected. However, an increase in vacuolization and the formation of large endosomes associated with peroxisomes were observed upon treatment (Fig. 6c). This cytological response coincided with a significant increase in the TMRM fluorescence signal in the intestinal lumen surrounding cells, indicating that the mitochondrial membrane potential was hyperpolarized (Fig. 6d-e). This phenomenon is crucial in cellular signaling and the regulation of diverse physiological processes, including mitochondrial ATP production, mitochondrial dynamics, and the induction of apoptosis. According to this response, transcriptome analysis of nematodes 24 h after treatment with cyclo(Pro-Tyr) revealed that 67 genes were induced and 381 genes were repressed (Fig. 6f). The inclusion of these genetic changes in metabolic pathways showed that fatty acid degradation, biosynthesis of unsaturated fatty acids or fatty acid elongation was repressed. In contrast, the double messenger system for signal transduction, phosphatidylinositol signaling system, calcium and MAPK signaling pathways were induced (Fig. 6g).

The most induced genes were as follows: i) Cytochrome P450, which is vital in maintaining cholesterol homeostasis in the cells. This gene family encodes a class of heme-containing enzymes that catalyze the oxidation of cholesterol, among other lipidic molecules. ii) Two genes encoding short-chain dehydrogenases that are known to catalyze the oxidation of various organic compounds, including alcohols, ketones, and aldehydes, to produce the corresponding acids or esters. iii) A CUB-like domain-containing protein, a motif found in extracellular and plasma membrane-associated proteins known to participate in tissue repair or toxic sensing. iv) Phospholipase A2 (PLA2) is responsible for the generation of free polyunsaturated fatty acids (PUFAs) by the degradation of phosphatidylcholine. v) Glutathione S-transferase, an enzyme that catalyzes the conjugation of reduced glutathione (GSH), which is an essential and polyvalent metabolite with special antioxidant functions; in addition, this enzyme is vital for cellular redox homoeostasis and development, growth or response to diverse stimuli. The most repressed genes included i) a PUFA elongation protein, which participates in reducing the amount of PUFAs, and ii) *pud-1.2*, which is abundantly expressed in the intestine and hypodermis and is indicative of this molecule target tissue (Fig. 6f).

## Discussion

The continuous interaction of microorganisms with cohabiting living cells relies on the secretion of a diverse battery of molecules, which can be specific or more generalist in their mode of action. *B. velezensis* is a soil-dwelling bacterium that can produce diverse bioactive molecules and potentially influence the nutritional network of the surrounding community^34^. In this work, we identified cyclo(Pro-Tyr) as an antifungal molecule and described how it exerts toxicity, which involves a mode of action also conserved against nematodes (Fig. 7). Although previously reported as antibacterial^9^, we have failed in reproducing such antibacterial activity against gram-positive or gram-negative bacteria (Supplementary Table 1, Fig. S8), which led us to support the specific reactivity of this molecule toward eukaryotic membranes.

**Figure 7.**
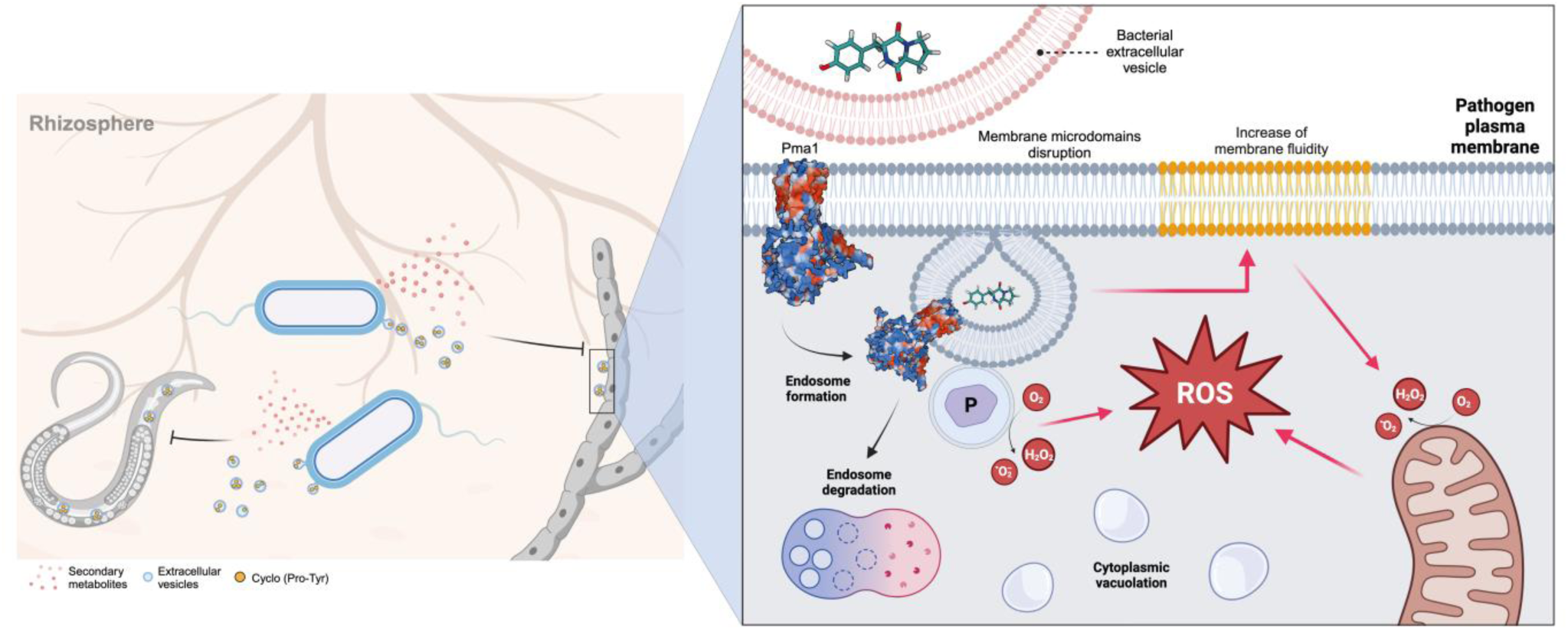
Overall scheme of the proposed mechanism for cyclo(Pro-Tyr). Left: In the dynamic environment of the rhizosphere, *B. velezensis* engages in intricate interactions with a variety of microorganisms, forging connections that influence the ecosystem’s delicate balance. This bacterium employs a multifaceted approach to interact with its neighboring organisms, which includes fungi and nematodes. One of the strategies employed by *B. velezensis* involves the secretion of bioactive small molecules, including cyclo(Pro-Tyr), most likely as part of vesicles. Right: Cyclo(Pro-Tyr) binds to Pma1, affecting folding or hexamerization. This impairment triggers a membrane damage alert signal, leading to the synthesis of ROS and prompting the removal of damaged membrane regions by endocytosis. The internalization and degradation of these regions, accompanied by the production of ROS, result in the dislocation of key proteins involved in membrane polarity maintenance, such as Pma1, and disruption of the organism’s homeostasis. Simultaneously, the decrease in phospholipids per unit volume increases membrane fluidity, becoming another source of stress contributing to the synthesis of ROS. Due to the increased membrane fluidity, the region is more permeable to that cyclodipeptides are released in vesicles, specialized carriers that can deliver bioactive molecules to their intended targets. Created with BioRender.com.

Through utilizing a sublethal concentration of cyclo(Pro-Tyr) (EC_50_), we monitored fungal adaptive responses and avoided rapid cellular collapse and death. The study offered a unique opportunity to uncover the complex dynamics between the cyclo(Pro-Tyr) compound and the target system. The two most noticeable fungal symptoms elicited by cyclo(Pro-Tyr) were the internalization of membrane sections via the formation of endosomes and activation of the oxidative burst (Fig. 7). Cellular components, such as proteins, lipids, and DNA, are targeted by ROS via oxidation and peroxidation^20,21^. Lipid peroxidation focuses on PUFAs, which involve hydrogen abstraction from a carbon; oxygen insertion produces lipid peroxyl radicals, hydroperoxides^23^ and lipid peroxides that can disrupt the integrity of cell membranes^35^. These chemical modifications can compromise the normal functioning of the plasma membrane and derived organelle membranes, affecting their permeability and stability^24^. Additionally, the accumulation of lipid peroxides can initiate a chain reaction, propagating further damage to neighboring lipids and amplifying the overall oxidative stress within the cell^20,22,23^. Diverse studies have reported a decrease in membrane fluidity in different cell membranes due to lipid peroxidation^24–26^; however, our findings suggest the opposite, i.e., membrane fluidity increases upon lipid peroxidation provoked by cyclo(Pro-Tyr). This discrepancy in the cellular response might be reflective of extreme inward membrane invagination and vesicle formation^27^, along with the segregation of cholesterol into distinct regions within the liquid crystalline membrane bilayer due to oxidative stress. The transfer of cholesterol from the phospholipid bilayer into separate cholesterol domains imposes a decrease in the overall width of the membrane bilayer, inducing an increase in fluidity in those cholesterol-impoverished regions^27–30^.

Most effects induced by cyclo(Pro-Tyr) are confined to the plasma membrane. Based on our findings, we proposed that the proton pump Pma1, which serves as a platform for organizing and regulating membrane-associated processes and is thus essential for cell viability, is the principal target of cyclo(Pro-Tyr). The most favorable proposed binding site for cyclo(Pro-Tyr) is situated in a domain involved in the hexamerization of Pma1^36^, suggesting that protein misfolding or arrest of hexamerization are the two complementary explanations for loss of protein functionality ^36,37^. Previous studies with *S. cerevisiae* have revealed that a misfolded version of Pma1 modified in the C-terminus is internalized in endosomes for further degradation in vacuoles. Additionally, this Pma1 mutant displays hypophosphorylation and fails to associate with Triton-insoluble fractions of the membrane, indicating a failure to enter lipid rafts^38^. Based on these studies and our experimental evidence, we propose that failure of Pma1 to oligomerize may trigger the internalization of membrane sections containing incorrectly folded proteins and, consequently, trigger the remaining physiological alterations associated with the disruption of membrane homeostasis. Thus, our findings support the rationale of using Pma1 as a target for designing novel antifungal molecules^39–41^. Methotrexate and fluconazole are two molecules that deplete Pma1 in *S. cerevisiae*, compromising ion transport and the influx of nutrients^29,42^. Plasma membrane disruption may render fungi more susceptible to the action of antifungal agents, overcoming resistance mechanisms and restoring the effectiveness of treatment. Demethylation inhibitors fungicides (DMI) disrupt ergosterol synthesis, and resistant fungal strains are known to overcome toxicity by restoring normal ergosterol levels within the plasma membrane^43,44^ using a variety of mechanisms: i) accumulating mutations in 14⍺-demethylase (CYP51), which reduce the affinity of DMIs to the target protein, ii) increasing the expression or copy number of the CYP51 gene, leading to heightened production of the target enzyme, or iii) overexpressing ATP-binding cassette (ABC) transporters that encode efflux pumps. According to our idea, we observed that the addition of cyclo(Pro-Tyr) to a myclobutanil-naturally resistant isolate of cucurbit powdery mildew, *Podosphaera xanthii*, heightened membrane permeability and resulted in the complete inhibition of fungal growth in the presence of myclobutanil (Fig. S9). Thus, we propose that the combined use of cyclo(Pro-Tyr) with existing DMIs antifungal drugs may lead to synergistic antifungal activity and a reduction in resistance development in fungal populations.

We observed that fungal lipid metabolism was rewired toward metabolites that should lead to increased membrane rigidity. One significant alteration involves the induction of phosphatidylserine decarboxylase, an enzyme that facilitates the conversion of phosphatidylserine into phosphatidylethanolamine, which is the second most abundant phospholipid of the membranes. Interestingly, the increase in level of phosphatidylethanolamine may increase the rigidity of the bilayer^24,45^. The following chemical variations also support the increase in *B. cinerea* membrane rigidity: i) the significant reduction in monoolein, an unsaturated lipid that significantly increases the fluidity of the lipid bilayer^46^, and ii) a reduction in the ergosterol byproduct ergocalciferol, which is described to increase membrane fluidity^47^. We also observed a reduction in palmitamide, an endogenous fatty acid synthesized from phosphatidylethanolamine at the plasma membrane^48^, and described it as a lipid messenger involved in cyclooxygenase activation for ROS detoxification^49^. A similar response in *C. elegans* was reproduced upon treatment with cyclo(Pro-Tyr), which elicited plasma membrane remodeling and enhanced the detoxification of ROS^50–52^. Furthermore, exposure of *C. elegans* to cyclo(Pro-Tyr) resulted in the upregulation of phospholipase A_2_ (PLA_2_), a crucial enzyme involved in the metabolism of membrane phospholipids; as a result PUFAs were produced, which are associated with a decrease in plasma membrane fluidity^27,53,54^.

The disruptions caused by cyclo(Pro-Tyr) can impact the interactions between organisms, nutrient cycling, and overall ecosystem functioning. The cycled chemical structure may be responsible for the activity and stability of cyclo(Pro-Tyr). However, the molecule exhibits a high hydrophobicity due to this definitory structural nature, which might impose difficulties in spreading the molecule in natural watery environments and diminish the ecological implications. We propose the secretion of the dipeptide within vesicles based on the following experimental evidence: i) mass spectrometry analysis revealed the presence of the molecule in isolated vesicles from cultures of *B. velezensis* WT and ii) the absence of the molecular ion in the vesicles produced by an attenuated vesicle forming the *xhlAB* mutant of *B. velezensis* (Fig. S10). Although not exclusive, we propose that the formation and secretion of vesicles is a general mechanism used by *Bacillus* and other bacterial species to manage the efficient delivery of these bioactive molecules in natural environments.

## Materials and methods

### Plant material

Melon (*Cucumis melo*; cv. Rochet Panal) and Zucchini (*Cucurbita pepo*; cv. Negro Belleza) plants were grown from seeds (Semillas Fito) at 25°C and 60% relative humidity under fluorescent and incandescent light at a photo fluency rate of ∼ 120 μmol m^-2^ s^-1^ and a 12/12 h photoperiod.

### Fungal strains—growth and inoculation of plant material

*B. cinerea* isolate B05.10 was cultured on potato dextrose agar (PDA, Oxoid) in a controlled-environment chamber at 20°C under illumination with fluorescent light at a photo fluency rate of 12 μmol m^-2^ s^-1^ and a 12/12 h photoperiod. Conidia were harvested from the light-grown culture in sterile distilled water containing 0.001% (v/v) Triton X-100 (J.T. Baker) and filtered through a 40-μm cell strainer to remove the remaining hyphae. Cell biology studies were carried out on germinated conidia grown in potato dextrose broth (PDB, Scharlau) inoculated with spore suspension incubated at 20°C for 24 h at 150 rpm.

*B. cinerea* plant inoculations were performed as previously described^1,55^. Briefly, the conidial suspension was adjusted to 10^5^ conidia mL^-1^ in half-strength filtered (0.45 μm) grape juice (100% pure organic). Melon plants 5-6 weeks old were used, and each leaf was inoculated with 5-μl droplets of conidial suspension (5·10^2^ conidia). Plants were covered with a plastic dome and placed in the growth chamber. At 72 h post-inoculation, leaves were imaged, and lesion size was analyzed using ImageJ 2.0 software. *Podosphaera xanthii* isolate 2208, which is resistant to DMI fungicides^56^, was grown on zucchini cotyledons, a cultivar that is very susceptible to powdery mildew, and maintained in vitro as previously described^57^.

A leaf-disc bioassay was used for fungicide-cyclodipeptide sensitivity testing. The conditions and methods necessary to carry out the fungicide sensitivity tests have been previously described^58^. Briefly, leaf discs from cotyledons of eight-day-old zucchini plants were placed adaxial surface down in petri dishes that contained sterile filter paper imbibed with 3 mL of 40 mg L^-1^ myclobutanil solution. After 24 h of incubation, the discs were transferred onto sterile filter paper that was deposited on Bertrand agar medium (sucrose 40 g L^-1^, benzimidazole 0.03 mL L^-1^, and agar 10 g L^-1^ in distilled water)^57^ in 5 cm diameter petri dishes, which remained open for approximately 1 h in a laminar airflow chamber at laboratory temperature to dry the discs. Afterward, leaf discs were treated with 0.25 mg mL^-1^ cyclo(Pro-Tyr) solution and air dried as described above. The discs were then inoculated on their adaxial surface with conidia of *P. xanthii* isolate 2208 with a disinfected eyelash. After 10 days of incubation, leaf discs were imaged, and the area covered by powdery mildew was calculated.

GFP-labeled *C. albicans* Pma1 was grown in YPD agar plates (1% peptone, 1% yeast extract, 2% glucose, and 2% agar) and pregrown in YPD medium (24 h, 28°C, 120 rpm). For experiments, 20 mL of YPD medium was inoculated with preculture, starting at A_600_= 0.1 and grown for 24 h (28°C, 120 rpm).

### Nematode strains and culture conditions

*C. elegans* wild-type N2, other mutants and transgenic worms were obtained from the Caenorhabditis Genetics Center (CGC) (see Supplementary Table 1). These strains were cultured at 20°C under standard growth conditions on NGM agar plates seeded with *E. coli* OP50. The population of *Meloidogyne incognita* was maintained on *Solanum lycopersicum* grown under greenhouse conditions.

For toxicity studies, the worms were synchronized using a bleaching solution (0.5 M NaOH and 1.2% NaClO). Then, the obtained embryos or L1 stage larvae were exposed to 0.25 mg mL^-1^ cyclo(Pro-Tyr) solution for 24 h at 20°C, and the effects of the chemical on survival rate, lifespan, and gene expression were measured. The Kaplan‒Meier survival curves were plotted by GraphPad Prism 9.0. For DIC and fluorescence microscopy, worms were placed on 3% agarose pads and mounted under a coverslip. DIC and fluorescence images were taken using a Zeiss LSM880 confocal laser scanning microscope equipped with a Plan-Apochromat 10x/0.45 M27 Oil objective. Images were false-colored by ImageJ.

For *C. elegans* synchronization, 4-day-old N2 plates were washed with 3 mL of M9 buffer and centrifuged at 1800 rpm for 1 min. The supernatant was removed carefully, and the worms were washed with M9 buffer at room temperature three to five times until the supernatant was clear. After the final centrifugation, the worm pellet was resuspended in bleaching solution (5 N NaOH, 5% NaClO) and vortexed for 3 min. Afterward, the eggs were pelleted and washed with M9 buffer. Finally, eggs were incubated at 20°C overnight with gentle shaking.

### Isolation of cyclo(Pro-Tyr) produced by *B. velezensis* CECT 8237

*B. velezensis* CECT 8237 was grown on lysogeny broth at 28°C and 150 rpm for 72 h. Afterward, the culture was centrifuged (10000 rpm, 10 min), and the filtered supernatant was extracted with butanol. After phase separation, the organic phase was concentrated to dryness at 60°C. The sediment was resuspended in methanol and subjected to SiO_2_ column chromatography using CHCl_3_/MeOH (9:1) as the mobile phase for fractionation. Fractions were collected to perform antimicrobial assays.

### Stress analysis of *B. cinerea* B05.10: identifying active fractions and critical concentration estimation

A stress analysis of *B. cinerea* B05.10 was performed to identify active fractions and estimate the critical concentration of purified cyclodipeptide. To identify active molecules, all fractions were incubated with *B. cinerea* germinated spores at 28°C for 24 h. Afterward, reactive oxygen species (ROS) production was analyzed using 10 mM hydrogen peroxide as a positive control and the solvent DMSO as a negative control. ROS detection was performed through dihydrorhodamine 123 (DHR123, Sigma‒ Aldrich). One microliter of DHR123 was added to a 1 mL cell suspension, followed by a 15 min incubation at room temperature in the dark. Images were obtained by visualizing samples using a Zeiss LSM880 confocal microscope with a Plan-apochromatic 63x/1.4 oil immersion objective and acquisitions with excitation at 488 nm. Image processing was performed using ImageJ software. All values were corrected for the corresponding image background. Once cyclo(Pro-Tyr) was identified, the critical concentration was calculated by incubation of a *B. cinerea* germinated spore suspension with 4, 2, 1, 0.5, 0.25, 0.125, 6.25·10^-2^, 3.13·10^-2^, 1.56·10^-2^, 7.81·10^-3^, 3.91·10^-3^, and 1.95·10^-3^ mg mL^-1^ Cyclo(Pro-Tyr) at 28°C and 150 rpm for 24 h, thus measuring the ROS response.

### Plasma membrane potential

To investigate the effect of cyclo(Pro-Tyr) treatment on the permeability of the plasma membrane, *B. cinerea* cells were incubated for 5 min with the voltage-sensitive fluorescent dye DiBAC_4_(3) (bis-(1,3-dibutylbarbituric acid)tri, methine oxonol; Thermo Fisher) at a final concentration of 20 μg mL^-1^. Images were obtained by visualizing samples using a Zeiss LSM880 confocal microscope with a Plan-apochromatic 63x/1.4 oil immersion objective and acquisitions with excitation at 488 nm. Cells were counterstained with lipophilic dye FM 4-64 (Thermo Fisher) to stain the plasma membrane.

### Plasma membrane fluidity

The effect of cyclo(Pro-Tyr) on plasma membrane fluidity was screened using two different methods. The first method was staining with DiIC_12_(3) dye (Thermo Fisher). Five milliliters of overnight *B. cinerea* culture was treated with 0.25 mg mL^-1^ Cyclo(Pro-Tyr) or DMSO as a control for 24 h min at 28°C and 150 rpm. Afterward, one *B. cinerea* clump was resuspended in 1 ml of 1× PBS, and 1 μg mL^-1^ DiIC_12_(3) was added. Cells were incubated at 28 °C for 30 min, washed three times and resuspended in 1× PBS. Images were obtained by visualizing samples using a Zeiss LSM880 confocal microscope with a Plan-apochromatic 63x/1.4 oil immersion objective and acquisitions with excitation at 561 nm. Image processing was performed using ImageJ software. For each experiment, the laser settings, scan speed, PMT detector gain, and pinhole aperture were kept constant for all acquired image stacks. The second method was fluorescence recovery after photobleaching (FRAP). For this procedure, 5 ml of overnight *B. cinerea* culture was treated with 0.25 mg mL^-1^ cyclo(Pro-Tyr) or DMSO as a control for 24 h min at 28°C and 150 rpm. One *B. cinerea* clump was placed onto a glass slide, and a reference image was taken. A central area of 2-3 μm was photobleached using a 405 nm laser at 80% output power, followed by immediate image acquisition at 5 sec intervals for 30 sec. Fluorescence recovery was measured as the average intensity in the bleached area of the plasma membrane. In parallel, the signal intensity in an unbleached area was measured. All fluorescent intensity values were corrected for adjacent image background, and signal intensities in photobleached regions were compared to those in unbleached parts of the plasma membrane.

### Mitochondrial membrane potential

Changes in mitochondrial membrane potential were determined with Image-iT™ TMRM Reagent (Thermo Fisher). One microliter of tetramethylrhodamine methyl ester (TMRM) was added to 1 ml of cell suspension, and the cells were further incubated at room temperature for 10 min in the dark and washed twice with PBS. For DIC and fluorescence microscopy, *C. elegans* N2 worms were placed on 3% agarose pads and mounted under a coverslip. Finally, stained cells were imaged using a 561 nm excitation wavelength. The intensity values were corrected by the cytoplasmic background.

### Lipid peroxidation

For microscopic analysis and quantification of lipid peroxidation in *B. cinerea* cells, the image-iT Lipid Peroxidation Kit (Thermo Fisher), based on BODIPY 581/591 C11 reagent, was used following the manufacturer’s instructions. Briefly, cyclodipeptide-treated *B. cinerea* cells were stained with a 10 µM solution of the image-iT Lipid Peroxidation sensor for 30 min. Finally, the cells were washed three times with PBS, mounted, and visualized immediately. The first image (oxidized channel) was acquired by exciting the sensor at 488 nm and recording the emissions at 510, followed by a second acquisition (reduced channel) with excitement at 561 nm and recording of the emissions at 590. The ratio of the emission fluorescence intensities at 590 nm to 510 nm gives the readout for lipid peroxidation in cells.

### Ergosterol measurement

The effect of cyclo(Pro-Tyr) on ergosterol content was assessed using filipin III staining (Sigma‒Aldrich). An overnight culture of *B. cinerea* B05.10 in potato dextrose broth (PDB) was incubated with 0.25 µg mL^-1^ cyclodipeptide at 28°C with agitation at 150 rpm for 24 h. Next, *B. cinerea* cells were stained with 50 µg mL^-1^ filipin dye for 2 min at room temperature. The images were visualized using a Zeiss LSM880 confocal microscope equipped with a Plan-apochromatic 63x/1.4 oil immersion objective, and acquisitions were performed with excitation at 405 nm.

### Transmission electron microscopy

*B. cinerea* and *C. elegans* were fixed in 2.5% (v/v) glutaraldehyde and 4% (v/v) paraformaldehyde in 0.1 M phosphate buffer (PB) overnight at 4°C. Worms were frozen in an HPM 100 high-pressure freezing machine (Leica Microsystems). Samples were postfixed in 1% osmium tetroxide solution in PB for 90 min at room temperature, followed by PB washes and 15 min of stepwise dehydration in an ethanol series (30%, 50%, 70%, 90% and 100% twice). Between the 50% and 70% steps, the samples were incubated in-bloc in 2% uranyl acetate solution in 50% ethanol at 4°C overnight. After dehydration, the samples were gradually embedded in low-viscosity Spurr’s resin (resin:ethanol, 1:1, 4 h; resin:ethanol, 3:1, 4 h; and pure resin overnight). The sample blocks were embedded in capsule molds containing pure resin for 72 h at 70°C. The samples were imaged under an FEI Tecnai G2 20 TWIN transmission electron microscope at an accelerating voltage of 80 kV. The images were acquired using TIA FEI Imaging Software v.4.14.

### Synthesis of cyclo(Pro-Tyr)

The synthesis of cyclodipeptide was carried out according to previously reported methods with minor modifications (Fig. S11). Briefly, Boc-Tyr-OH (563 mg, 2.0 mmol) was dissolved in anhydrous CH_2_Cl_2_ (10 mL) at 0°C. To this mixture, the following reactants were consecutively added: H-Pro-OMe•HCl (331 mg, 2.0 mmol), Et_3_N (0.28 mL, 2.0 mmol) and EDC•HCl (384 mg, 2.0 mmol). After 16 h of stirring at 4°C, the mixture was washed with a citric acid solution (1 M) followed by a saturated solution of NaHCO_3_. The organic phase was dried with anhydrous MgSO_4_, filtered and evaporated under reduced pressure. The resulting white solid (**1**) was dissolved without further purification in a minimal amount of anhydrous CH_2_Cl_2_ at 0°C, and anhydrous MeOH (0.2 mL) was added to the mixture. Then, acetyl chloride (0.28 mL, ∼4 equiv.) was added dropwise to the reaction. After 5 h of stirring at 0°C, the solvent was removed on a rotary evaporator, affording a pale-yellow oil (**2**) that was dried on high vacuum overnight. Cyclo(Pro-Tyr) (**3**) was obtained after **2** was dissolved in anhydrous DMF (2 mL) at room temperature and cyclized and piperidine was added (0.2 mL, ∼3 equiv.). The resulting mixture was concentrated in a rotary evaporator to afford a pale-yellow oil that was washed with citric acid (1 M), dried with anhydrous MgSO_4_, filtered, and evaporated to dryness. Further purification was performed on a silica gel column with MeOH/CH_2_Cl_2_ (1:9) as the mobile phase to yield 94 mg of a white powder (**3**, 24% overall yield). Compounds were confirmed using a 500 MHz Bruker 500 Advance III NMR spectrometer. A proton experiment was run to confirm the structure. MALDI-TOF-TOF mass spectra were recorded on a Bruker Ultraflex Extreme system (Bruker Daltonics, Billerica, MA, USA). ^1^H-NMR: (400 MHz, MeOD) δ 7.04 (d, 2H, *J*=7,0 Hz, H-12 and H-16), 6,81 (d, 2H, *J*=6,7 Hz, H-13 and H-15), 4,35 (t, 1H, *J*=2.8 Hz, H-3), 4,04 (ddd, 1H, *J*=1.2 and 1,6 Hz, H-6), 3,54 (dt, 1H, *J*=9,2 and 6,8 Hz, H-9a), 3,35 (M, 1H, H-9b), 3,06 (qd, 2H, *J*= 4,0 and 4,4 Hz, H-10a and H-10b), 2,09 (m, 1H, H-8a); 1,80 (m, 2H, H-8b and H-7a), 1,23 (m, 1H, H-7b). Mass spectrum (MS/MS-TOF), *m/z*: 262 [M+H^+^]^+^, 243 [M-CO+H^+^]^+^, 232 [M-CO-NH_3_+H^+^]^+^, 217 [M-2CO^+^]^+^, 146 [M-residue mass of Pro-NH_3_+H^+^]^+^, 135 (Immonium ion of Tyr), 106 Hydroxybenzyl ion from Tyr), 97 [M-residue mass Tyr+H^+^]^+^, 70 (immonium ion of Pro). HRMS: 261.1234 [M^+^+H^+^]^+^ (Theoretical: 260.116).

### Synthesis of a fluorophore-derivatized cyclo(Pro-Tyr)

Over a solution of **3** (45 mg, 0.17 mmol) in anhydrous CH_2_Cl_2_ (4 mL) and anhydrous pyridine (132 μL), DCC (146 mg, 0.71 mmol) and fluorescein sodium salt (25 mg, 0.06 mmol) were added at 0°C and stirred at this temperature for 2 h and then at room temperature for an additional 2 h. The reaction was finished by evaporating the solvent under N_2_ stream, and the residue was dissolved in MeOH (HPLC quality) to be further separated on a preparative HPLC (Agilent infinity 1260, LC column Luna® 5μm C18, 100 Å, 150 x 10 mm; mobile phase: A = Acetonitrile B = 0.1% HCOOH, Flow = 2 mL min^-1^, λ = 254 nm).

### Immunofluorescence microscopy

*B. cinerea* cells incubated with 0.25 mg mL^-1^ cyclo(Pro-Tyr) for 24 h were fixed with fixation solution (0.1% (v/v) glutaraldehyde and 3% (v/v) paraformaldehyde in 1× PBS) for 15 min at room temperature. After being washed with 1× PBS, the cells were blocked with blocking solution (3% (w/v) BSA and 0.2% (v/v) Triton X-100) for 60 min. Finally, the cells were stained for immunofluorescence using a 1:50 mouse monoclonal anti-Pma1 primary antibody (Thermo Fisher) followed by a 1:200 GFP-conjugated rabbit anti-mouse secondary antibody (Jackson ImmunoResearch). Images were obtained by visualizing samples using a Zeiss LSM880 confocal microscope with a Plan-apochromatic 63x/1.4 oil immersion objective and acquisitions with excitation at 488 nm. Cells were counterstained with the lipophilic dye FM 4-64 (Thermo Fisher) to stain early endosomes.

### Isolation of DRM from *B. cinerea* cells

DRM was isolated as described previously^29^, with modifications. *B. cinerea* was grown for 24 h in PDB medium; then, clumps were pelleted and washed twice in 0.9% NaCl. Cells were resuspended in 1 mL of TNE buffer (50 mM Tris pH 7.5, 150 mM NaCl, 5 mM EDTA), followed by the addition of the same volume of 1 mm tungsten carbide beads. Cells were broken by a bead beater, beads were removed, and the lysate was cleared by centrifugation (5000 rpm, 3 min, 4°C). The lysate was transferred to a new tube, with the addition of Triton X-100 to a final concentration of 3% and incubated for 30 min on ice.

A total of 10 mL of sample was pipetted carefully into previously prepared ultracentrifuge tubes. Samples were ultracentrifuged at 35,000 rpm for 1.5 h at 4°C. Afterward, the pellet was resuspended in 1× PBS, and the total protein content was precipitated with TCA to a final concentration of 10% and incubated on ice for 1 h. Samples were centrifuged (12,000 rpm, 15 min, 4°C), and the pellet was washed twice with cold acetone. After acetone was completely evaporated, the pellet was resuspended in 20 mM Tris-HCl (pH = 8.0) and 20 mM NaCl. For electrophoresis, 15 µL was used and resuspended in 2x 𝛽-mercaptoethanol loading buffer, run on a 12% SDS‒PAGE gel and transferred onto a PVDF membrane using a Trans-Blot Turbo electrophoretic transfer cell (Bio-Rad) at 25 V for 30 min. The blot was probed with 1:1000 anti-Pma1 mouse polyclonal antibody (Thermo Fisher). A secondary anti-mouse antibody conjugated to horseradish peroxidase (Bio-Rad) was used at a dilution of 1:20000. The blot was developed using the Pierce super signal detection system following the supplier’s recommended procedures (Pierce).

### Biofilm assays

Biofilm formation of *Bacillus* species was analyzed using MSgg medium (100 mM morpholinopropanesulfonic acid (MOPS) (pH 7), 0.5% glycerol, 0.5% glutamate, 5 mM potassium phosphate (pH 7), 50 μg mL^-1^ tryptophan, 50 μg mL^-1^ phenylalanine, 2 mM MgCl_2_, 700 μM CaCl_2_, 50 μM FeCl_3_, 50 μM MnCl_2_, 2 μM thiamine, and 1 μM ZnCl_2_), as previously described^59^. For precultures, each strain was cultivated on LB agar supplemented with the appropriate antibiotic at 37°C for 8 h. A single colony was then resuspended in 1 mL of MSgg medium, and 10 μL of this suspension was used to inoculate 1 mL of MSgg in 24-well plates supplemented with 0.25 mg mL^-1^ cyclo(Pro-Tyr) or DMSO as a solvent control. The plates were then incubated at 30°C without agitation. The presence of wrinkles, a morphological characteristic of mature *Bacillus* biofilms, was examined in the pellicles formed during the incubation.

The biofilm formation of gram-negative bacteria was assessed using bacterial adhesion to abiotic surfaces. Cultures were grown in LB medium at 30°C without agitation overnight in the presence of 0.25 mg mL^-1^ cyclo(Pro-Tyr) or DMSO as a solvent control. To initiate the staining process, 1 mL of a 1% solution of crystal violet in water was added to each well of a 24-well plate. After a 5-min incubation, the plates were rinsed five times by immersion in tap water and then left inverted to dry on the bench for at least 45 min. Subsequently, the crystal violet was resuspended using 50% acetic acid, and the resuspended solution was further diluted at a ratio of 1/10. The absorbance of the diluted solution was measured at 595 nm to quantify the extent of biofilm formation.

### Protein modeling and molecular docking

AlphaFold^60^ was used for automated protein tertiary structure modeling of *Botrytis cinerea* B05.10 Pma1 protein. To identify potential binding sites of cyclo(Pro-Tyr) (PubChem ID: 119404) to the Pma1 protein, automated molecular docking and thermodynamic analysis were performed using the web-based “SwissDock program [www.swissdock.ch/docking]”^61^. SwissDock predicts the possible molecular interactions between a target protein and small molecule based on the docking algorithm EADock DSS^62^. The docking was performed using the “Accurate” parameter at otherwise default parameters, with no region of interest defined (blind docking). Binding energies are estimated by using CHARMM (Chemistry at HARvard Macromolecular Mechanics), a molecular simulation program implemented within SwissDock software, and the most favorable energies are evaluated by FACTS (Fast Analytical Continuum Treatment of Solvation). Finally, the energy results are scored and ranked by *full fitness* (kcal mol^−1^), and spontaneous binding is exhibited by the estimated Gibbs free energy ΔG (kcal mol^−1^). The negative values of ΔG support the assertion that the binding process is highly spontaneous. Modeling and docking results were visualized using UCSF Chimera v1.8 software.

### Sample preparation for NMR

Four samples were prepared. For each mixture, appropriate amounts of lipid (POPC, POPC-D31, and POPE) with and without the dipeptide were co-solubilized in chloroform. Chloroform was evaporated under N_2_ flux to obtain a lipid film. Then, 1 mL of water was added and each mixture was lyophilized overnight. 100 μL of depleted (2-3ppm ^2^H content) water was added. Next, the samples were subjected to three thermal cycles, which involved placing them at -198°C in liquid N2 for a few seconds and then in a water bath at 40°C for 10min. The samples were transferred to 4mm NMR rotors.

### Solid-state NMR spectroscopy

Solid state ^2^H-NMR spectra were acquired by means of a quadrupolar echo pulse sequence (90°- τ- 90°- τ- acq) on an Avance III WB 500 NMR spectrometer (Bruker Biospin, Wissembourg, France) operating at 500MHz for ^1^H and 76.7MHz for ^2^H equipped with a dual 1H/X 4-mm MAS probe. The acquisition parameters were as follow: spectral window of 500 KHz, π/2 pulse width of 3.5μs, recycle delay of 2s, and echo delay of 50 μs, number of scans 512. Samples were allowed to equilibrate for 20 min at a given temperature before the NMR signal was acquired. Experimental temperatures were from 268K to 318K every 5K and an additional step at 310K. Lorentzian noise filtering with a width of 500Hz was applied prior Fourier transformation from the top of echo signal.

Spectral simulations: A user-friendly software interface was developed by Arnaud Grélard using computer programs written in FORTRAN by Erick Dufourc: the time domain trace composed of the weighed sum of signals corresponding to each quadrupolar splitting (most intense doublet of the Pake pattern) of individual C-D bonds of the entire lipid chain is calculated, and a Fourier transformation is performed, leading to spectra as in Figure S1. Adjustable parameters are the individual quadrupolar splittings and their intrinsic line width. All measurable quadrupolar splittings are taken as input parameters, the weight of each splitting being set according to the molecular structure. Comparison between experimental spectra and calculated is made until a satisfactory superimposition is obtained.

Calculation of first spectral moments: First moments were calculated using a python home-made routine (Sébastien Buchoux, NMRDepaker) to which Bruker NMR data were exported.

### RNA isolation

RNA from *B. cinerea* B05.10 was extracted from clumps after 24 h of growth on PDB supplemented with 0.25 mg mL^-1^ cyclo(Pro-Tyr) and DMSO as a control at 28°C and 150 rpm using a NucleoSpin RNA Plant and Fungi kit (Macherey-Nagel) according to the manufacturer’s instructions.

RNA from synchronized *C. elegans* N2 was extracted as previously described^7^ with slight modifications. Briefly, *C. elegans* N2 was incubated in M9 buffer supplemented with 0.25 mg mL^-1^ cyclo(Pro-Tyr) and DMSO as a control for 24 h at 19°C with gentle shaking. Afterward, worms were centrifuged, and the supernatant was discarded. The worm pellet was resuspended in TRI-Reagent (Simga-Aldrich) with 0.1 mm glass beads, and worms were lysed in TissueLyser II (Qiagen) at 30 cycles sec^-1^ for 20 min. After worm lysis, RNA was extracted using the NucleoSpin RNA Plant and Fungi kit (Macherey-Nagel) according to the manufacturer’s instructions.

### Transcriptome analysis

For RNA sequencing analysis, 200-500 bp paired-end read libraries were prepared using a TruSeq stranded total RNA kit (Illumina). The libraries were sequenced using a NextSeq550 instrument (Illumina). Raw reads were preprocessed by NextSeq System Suite v.2.2.0. using specific NGS technology configuration parameters^63^. This preprocessing removes low-quality, ambiguous and low-complexity stretches, linkers, adapters, vector fragments and contaminated sequences and preserves the longest informative parts of the reads. SeqTrimNext also discarded sequences shorter than 25 bp. Subsequently, clean reads of the BAM files were aligned and annotated by Bowtie2 using the *B. cinerea* B05.10 v.109.3^64^ and *C. elegans* N2^65^ genomes as references; these data were then sorted and indexed using SAMtools v.1.484110. Uniquely localized reads were used to calculate the read number values for each gene by Sam2counts (https://github.com/vsbuffalo/sam2counts). Differentially expressed genes (DEGs) in the treatment samples were analyzed by DEgenes Hunter, which provides a combined *P* value calculated based on the Fisher method using nominal *P* values provided by edgeR and DEseq2. This combined *P* value was adjusted by the Benjamini‒Hochberg procedure (false discovery rate approach) and used to rank all obtained DEGs. For each gene, a combined *P* value < 0.05 and log_2_-fold change (FC) >1 or <−1 were considered significance thresholds. A heatmap and DEG clustering were generated using ComplexHeatmap in R Studio and Kobas 2.0. Only profiles with a *P* value < 0.05 were considered in the present study. DEGs annotated using the *B. cinerea* genome were processed to identify the Gene Ontology functional categories using sma3s and TopGo software. Kyoto Encyclopedia of Genes and Genomes (KEGG) pathways and enrichment were estimated using Bioconductor packages (Bioconductor.org) GGplot2, ClusterProfiler, DOSE and EnrichPlot in R Studio. The data were deposited in the GEO database under the reference GSE237225 and GSE237224 for *B. cinerea* B05.10 and *C. elegans* N2, respectively.

### Metabolite extraction

Metabolites from the samples of *B. cinerea* treated with cyclo(Pro-Tyr) and untreated were extracted by adding 1 mL methanol to a suspension of *B. cinerea* clumps grown in PDB overnight at 28°C and disrupting the tissue in TissueLyser at 30 cycles s^-1^ for 30 min with 1 mm tungsten carbide beads. Two replicates were performed for each experiment.

Differences in the abundance of metabolites were estimated on the basis of relative comparisons between equally treated samples due to the absence of authentic standards in the nontargeted metabolomics approach.

### Liquid chromatography-tandem mass spectrometry (LC‒MS/MS)

Nontarget metabolomics was performed by micro-flow liquid chromatography-tandem mass spectrometry (LC‒MS/MS) with a UHPLC coupled to a Q Exactive HF mass spectrometer and optimized data-dependent acquisition method as previously described^66^. In brief, UHPLC separation was performed using a C18 core-shell column (Kinetex, 50 × 1 mm, 1.7 µm particle size, 100 A pore size, Phenomenex, Torrance, USA). The mobile phases used were solvent (A) H2O (LC/MS grade, Fisher Scientific) + 0.1% formic acid (FA) and solvent (B) acetonitrile (LC/MS grade, Fisher Scientific) + 0.1% FA. After the sample injection, a linear gradient method of 5 min was used for elution of small molecules, and the flow rate was set to 150 µL min^-1^ (microflow). The following separation conditions were set as follows: in the time range 0-4 min from 5% to 50% solvent (B) was used, 4-5 min from 50 to 99% B, followed by 2 min washout phase at 99% B and 3 min re-equilibration phase at 5% B. The measurements were conducted in positive mode, and the heated electrospray ionization (HESI) parameters included a sheath gas flow rate of 30 AU, auxiliary gas flow rate of 10 AU, and sweep gas flow rate of 2 AU. The spray voltage was set to 3.50 kV, the inlet capillary temperature was set to 250°C, the S-lens RF level was set to 50 V, and the auxiliary gas heater temperature was set to 200°C. The full MS survey scan acquisition range was set to 120–1,800 m/z with a resolution of 45,000, automatic gain control (AGC) of 1E6, and maximum injection time of 100 ms with one microscan. MS/MS spectra acquisition was performed in data-dependent acquisition (DDA) mode with TopN set to 5; as a consequence, the five most abundant precursor ions of the survey MS scan were used for MS/MS fragmentation. The resolution of the MS/MS spectra was set to 15,000, the AGC target to 5E5 and the maximum injection time to 50 ms. The quadrupole precursor selection width was set to 1 m/z. Normalized collision energy was applied stepwise at 25, 35, and 45. MS/MS scans were triggered with apex mode within 2–15 s from their first occurrence in a survey scan. Dynamic precursor exclusion was set to 5 s.

### Data analysis and MS/MS network analysis

After LC‒MS/MS acquisition, raw spectra were converted to mzML files using MSconvert (ProteoWizard). MS1 and MS/MS feature extraction was performed with Mzmine3^67^. For MS1 spectra, an intensity threshold of 1E5 was used, and for MS/MS spectra, an intensity threshold of 1E3 was used. For MS1 chromatogram building, a 10-ppm mass accuracy and a minimum peak intensity of 5E5 were set. Extracted ion chromatograms (XICs) were deconvolved using the baseline cutoff algorithm at an intensity of 1E5. After chromatographic deconvolution, XICs were matched to MS/MS spectra within 0.02 m/z and 0.2-min retention time windows. Isotope peaks were grouped, and features from different samples were aligned with 10 ppm mass tolerance and 0.1-min retention time tolerance. MS1 features without MS2 features assigned were filtered out of the resulting matrix as well as features that did not contain isotope peaks and that did not occur in at least three samples. After filtering, gaps in the feature matrix were filled with relaxed retention time tolerance at 0.2 min but also 10 ppm mass tolerance. Finally, the feature table was exported as a.csv file, and the corresponding MS/MS spectra were exported as .mgf files. Contaminate features observed in blank samples were filtered, and only those with a relative abundance ratio blank to average lower than 30% were considered for further analysis. For feature-based molecular networking and spectrum library matching, the .mgf file was uploaded to GNPS (https://gnps.ucsd.edu/ProteoSAFe/static/gnps-splash.jsp)^68^.

For molecular networking, the minimum cosine score was set to 0.7. The precursor ion mass tolerance was set to 0.01 Da, and the fragment ion mass tolerance was set to 0.01 Da. The minimum number of matched fragment peaks was set to 6, the minimum cluster size was set to 1 (MS cluster-off), and the library search minimum number of matched fragment peaks was set to 6. When analog searches were performed, the cosine score threshold was 0.7, and the maximum analog search mass difference was 100 m/z. Molecular networks were visualized with Cytoscape version 3.9.1^69^.

To enhance the chemical structural information in the molecular network, the generated mgf file from MZmine3 was placed into SIRIUS 4 for chemical class and structure prediction. The input data were automatically compared and classified against databases present in SIRIUS (Bio Database, GNPS, Natural Products, PubChem, PubMed). For Molecular Formula Identification, MS2 mass accuracy was set to 3 ppm. Chemical class annotations were performed with CSI: FingerID and CANOPUS using ClassyFire chemical ontology.

Mirror plots were inspected through the spectrum resolver at GNPS (https://metabolomics-usi.ucsd.edu/), comparing mzspec of the selected features and the metabolites recorded in MS/MS databases (Fig. S12). Statistical analysis of metabolomic data sets was performed by Metaboanalyst v.5.0 after data filtering by interquartile range (IQR).

### Statistical analysis

When data were normally distributed and sample variances were equal, t-tests were performed. In all other cases, the Mann‒Whitney Rank Sum test was performed. For multiple comparisons, one-way analysis of variance (ANOVA) was performed when the equal variance test was passed. In all other cases, one-way ANOVA on ranks was performed (Kruskal‒Wallis significant difference test). Significance was accepted at P < 0.05.

## Acknowledgments

We thank Saray Morales Rojas for its technical support, Alicia Esteban and David Navas from the IHSM and SCAI microscopy units, respectively, for their technical support in confocal microscopy, Josefa Gómez Maldonado from the Ultrasequencing Unit of the SCBI-UMA for RNA sequencing, Luis Díaz and Belén Delgado-Martín from the Bioinformatic Unit of UMA and Mercedes Martín Rufián from the Proteomic Unit of the SCAI-UMA for technical suggestions and LC‒MS analysis, and Manuel Muñoz and Ana María Brokate from University Pablo de Olavide for supplying nematodes and technical support. This work was partially supported by grants from ERC Starting Grant (BacBio 637971), Plan Nacional de I+D+I of Ministerio de Economía y Competitividad (PID2019-107724GB-I00, PID2022-141664NB-I00), research contract 8.06/60.4086 with KOPPERT B.V. (The Netherlands), and Proyecto Jóvenes Investigadores from Plan Propio de Universidad de Málaga (B1-2021_34). D.V.C. is funded by the program Incorporación de Doctores PAIDI from Junta de Andalucía (DOC_00266). A.I.P. L is funded by the program FPU (FPU19/00289), JAE-Intro 2019 (JAEINT 19 00269) and the program Plan Propio de Investigación y Transferencia from Universidad de Málaga. D.P. was supported by the Deutsche Forschungsgemeinschaft through the CMFI Cluster of Excellence (EXC 2124) and the Collaborative Research Center CellMap (TRR 261). P.S. is supported by the European Union’s Horizon Europe research and innovation programme through a Marie Skłodowska-Curie fellowship n.101108450-MeStaLeM.

## Author contributions

D.R. conceived the study; D.R. and D.V.C. designed the experiments; D.V.C. performed the main experimental work; D.V.C. performed and designed the confocal microscopy work and data analysis; D.V.C. performed docking analyses; D.V.C. performed RNAseq and genomic data analyses; J.H. performed cyclodipeptide synthesis and cyclodipeptide labeling. A.I.P. L, P.S. A.K.P.S. and D.P. performed MALDI-TOF MSI and LC‒MS/MS analysis; A.G. and A.L. performed all biophysical analyses; D.R. and D.V.C. wrote the manuscript; and A.V., and A.P.G. contributed critically to writing the final version of the manuscript.

---

**Supplementary table 1.** Minimum inhibitory concentration of cyclo(Pro-Tyr) on Bacteria.

| Bacterial Strains | MIC (mg mL<sup>-1</sup>) |
| --- | --- |
| *Bacillus velezensis* CECT 8238 | >10 |
| *Bacillus velezensis* FZB42 | >10 |
| *Bacillus subtilis* 3610 | >10 |
| *Bacillus subtilis* 168 | >10 |
| *Pseudomonas putida* KT2440 | >10 |
| *Pseudomonas aeruginosa* PAO1 | >10 |
| *Aeromonas hydrophila* ATCC 7966 | >10 |
| *Erwinia persicina* ATCC 35998 | >10 |

**Supplementary table 2.** C. *elegans* lines used in the present study.

| Strain name | Genotype | Description |
| --- | --- | --- |
| FT63 | *xnls17* | *xnls17* [*dlg-1::GFP* + *rol-6*(su1006)]. Rollers. Expresses DLG-1::GFP in epithelial cell junctions. |
| CL2122 | *dvls15* | *dvls15* [(pPD3038) *unc54* (vector) + (pCL26) *mtl-2::GFP*]. Control strain for CL2120. Phenotype apparently WT. |
| BOX213 | *erm-1*(*mib15*[*erm-1::eGFP*]) | Endogenous *erm-1* locus tagged with eGFP. Homozygous viable, partially functional endogenous *erm-1* tag. *erm-1::GFP* animals have a reduced brood size and incomplete outgrowth of the excretory canal, but show no other developmental or morphological abnormalities. |

