## Supplementary material for "Cyclo(Pro-Tyr) elicits conserved cellular damage in fungi by targeting the [H^+^]ATPase Pma1 in plasma membrane domains": Suplementary figures: Suplementary figures.pptx

### Slide 1
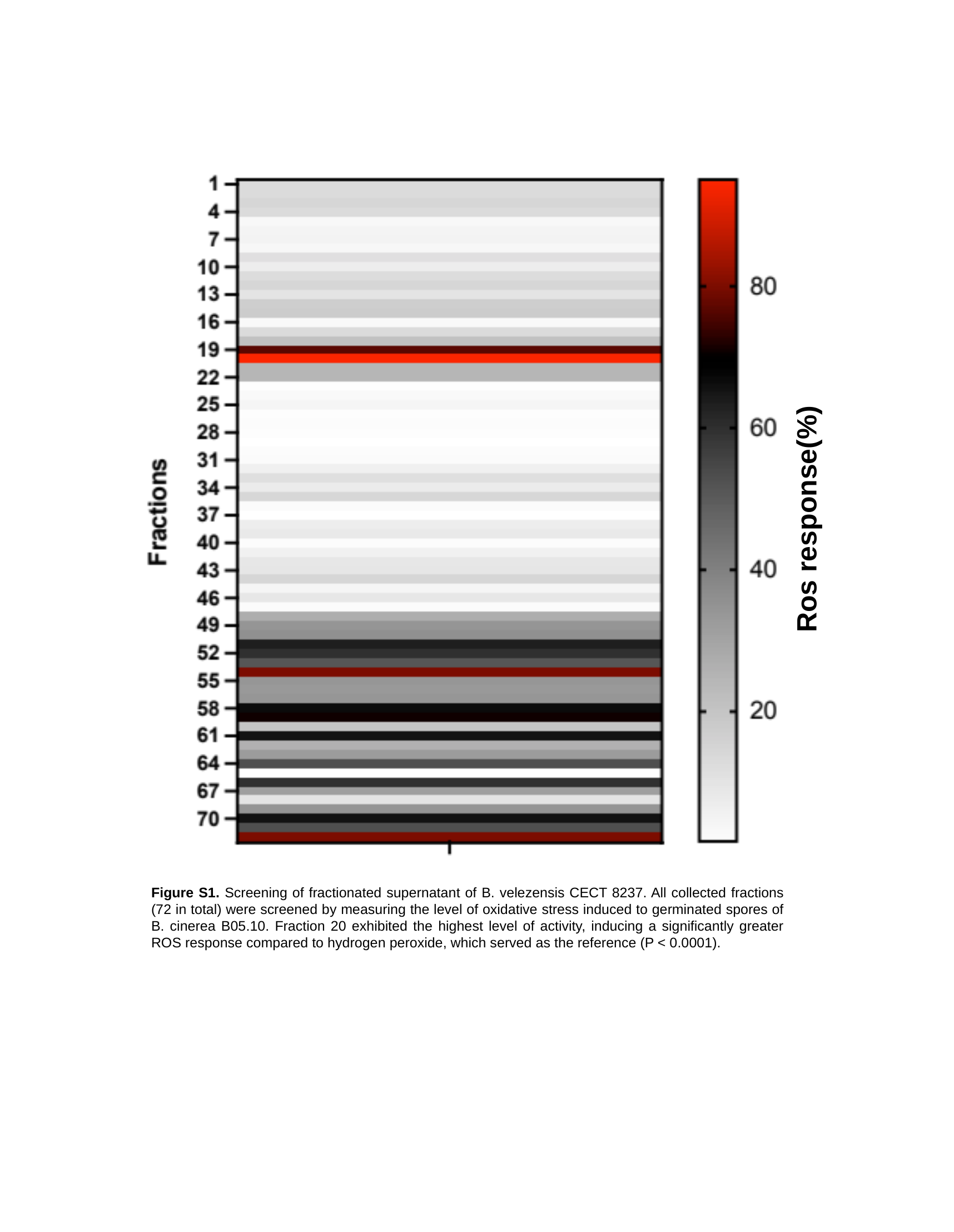

Ros response(%)
Figure S1. Screening of fractionated supernatant of B. velezensis CECT 8237. All collected fractions (72 in total) were screened by measuring the level of oxidative stress induced to germinated spores of B. cinerea B05.10. Fraction 20 exhibited the highest level of activity, inducing a significantly greater ROS response compared to hydrogen peroxide, which served as the reference (P < 0.0001).

### Slide 2
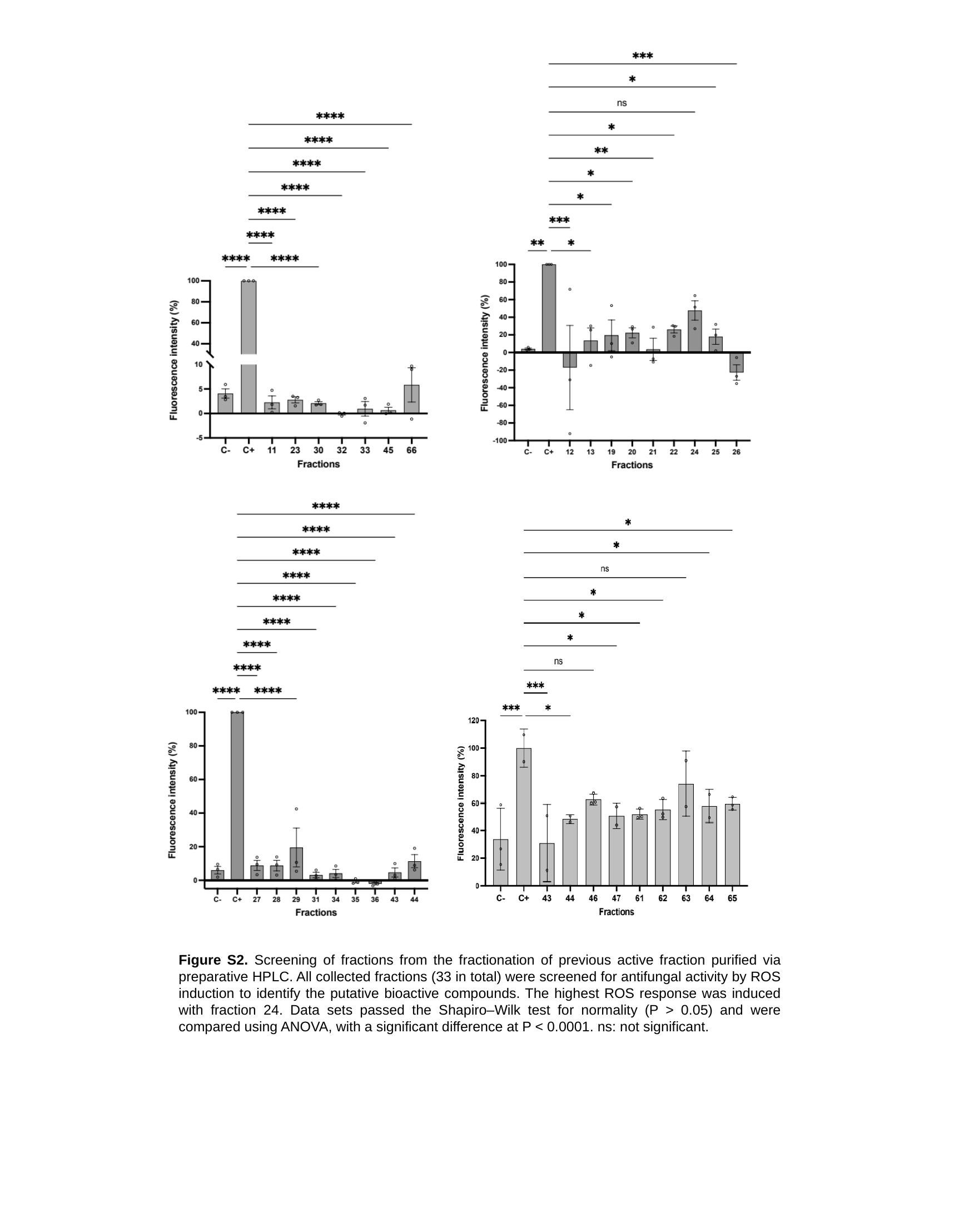

Figure S2. Screening of fractions from the fractionation of previous active fraction purified via preparative HPLC. All collected fractions (33 in total) were screened for antifungal activity by ROS induction to identify the putative bioactive compounds. The highest ROS response was induced with fraction 24. Data sets passed the Shapiro‒Wilk test for normality (P > 0.05) and were compared using ANOVA, with a significant difference at P < 0.0001. ns: not significant.

### Slide 3
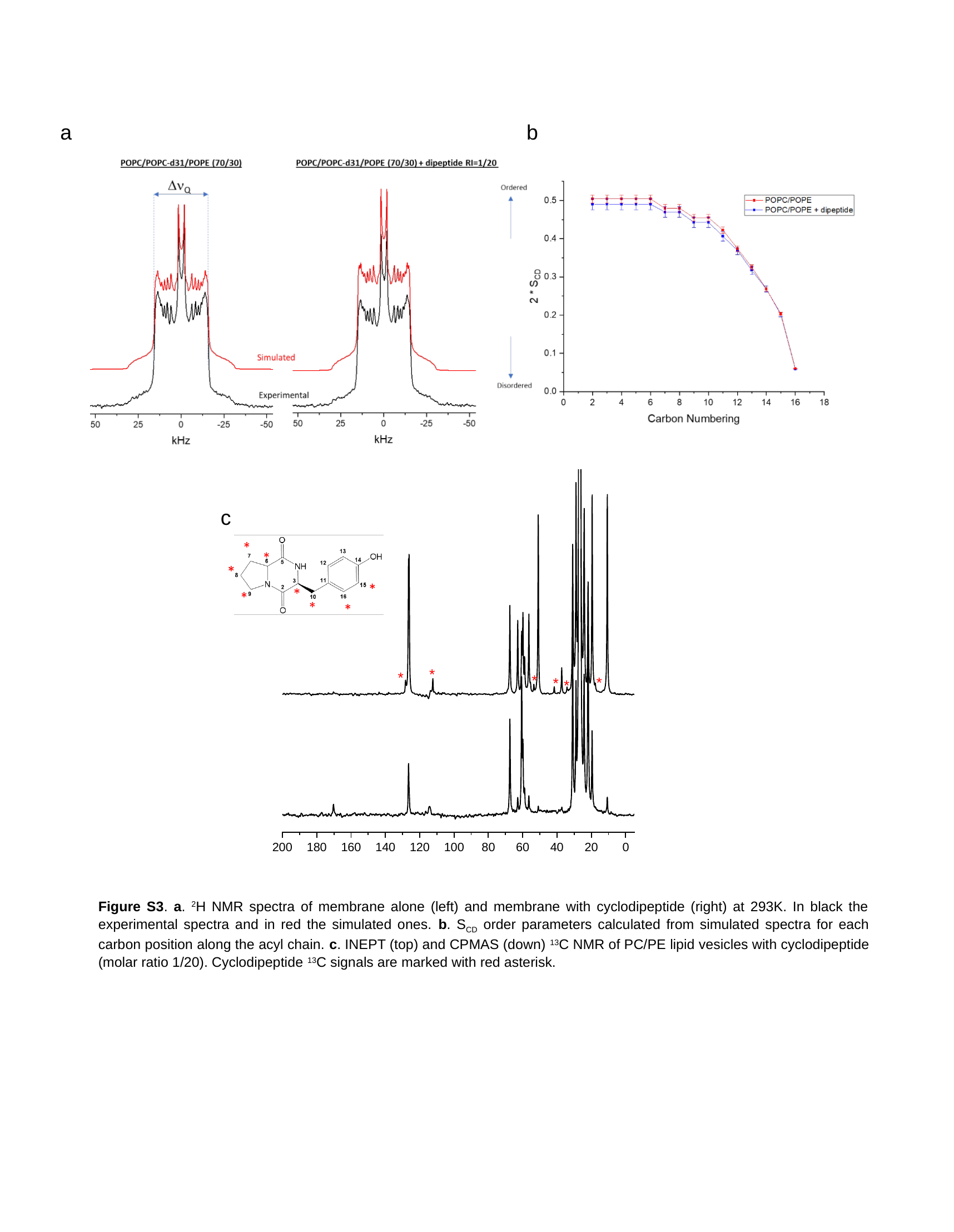

a
b
c
Figure S3. a. ^2^H NMR spectra of membrane alone (left) and membrane with cyclodipeptide (right) at 293K. In black the experimental spectra and in red the simulated ones. b. S_CD_ order parameters calculated from simulated spectra for each carbon position along the acyl chain. c. INEPT (top) and CPMAS (down) ^13^C NMR of PC/PE lipid vesicles with cyclodipeptide (molar ratio 1/20). Cyclodipeptide ^13^C signals are marked with red asterisk.

### Slide 4
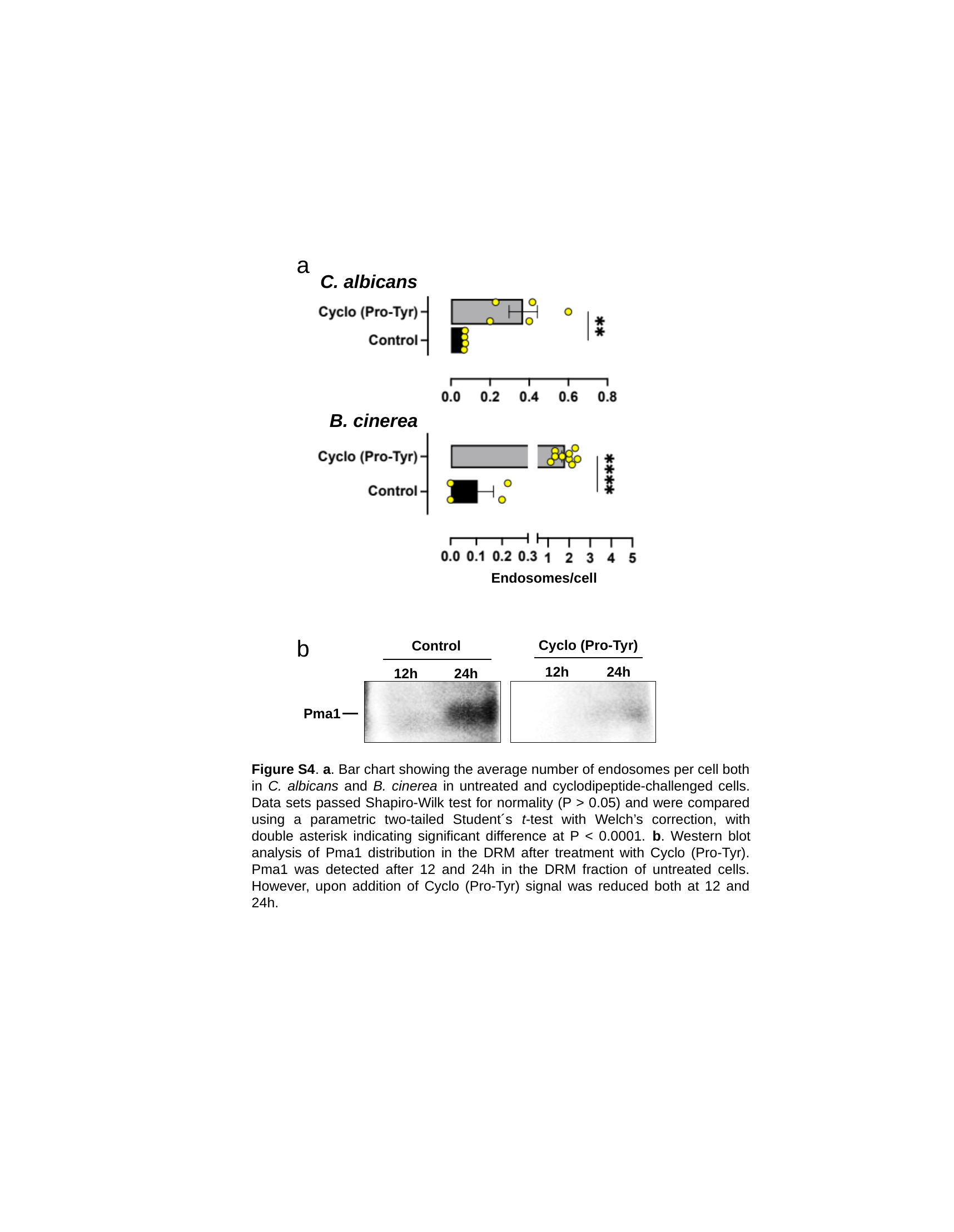

a
C. albicans
B. cinerea
Endosomes/cell
b
Cyclo (Pro-Tyr)
12h
24h
Control
12h
24h
Pma1
Figure S4. a. Bar chart showing the average number of endosomes per cell both in C. albicans and B. cinerea in untreated and cyclodipeptide-challenged cells. Data sets passed Shapiro-Wilk test for normality (P > 0.05) and were compared using a parametric two-tailed Student´s t-test with Welch’s correction, with double asterisk indicating significant difference at P < 0.0001. b. Western blot analysis of Pma1 distribution in the DRM after treatment with Cyclo (Pro-Tyr). Pma1 was detected after 12 and 24h in the DRM fraction of untreated cells. However, upon addition of Cyclo (Pro-Tyr) signal was reduced both at 12 and 24h.

### Slide 5
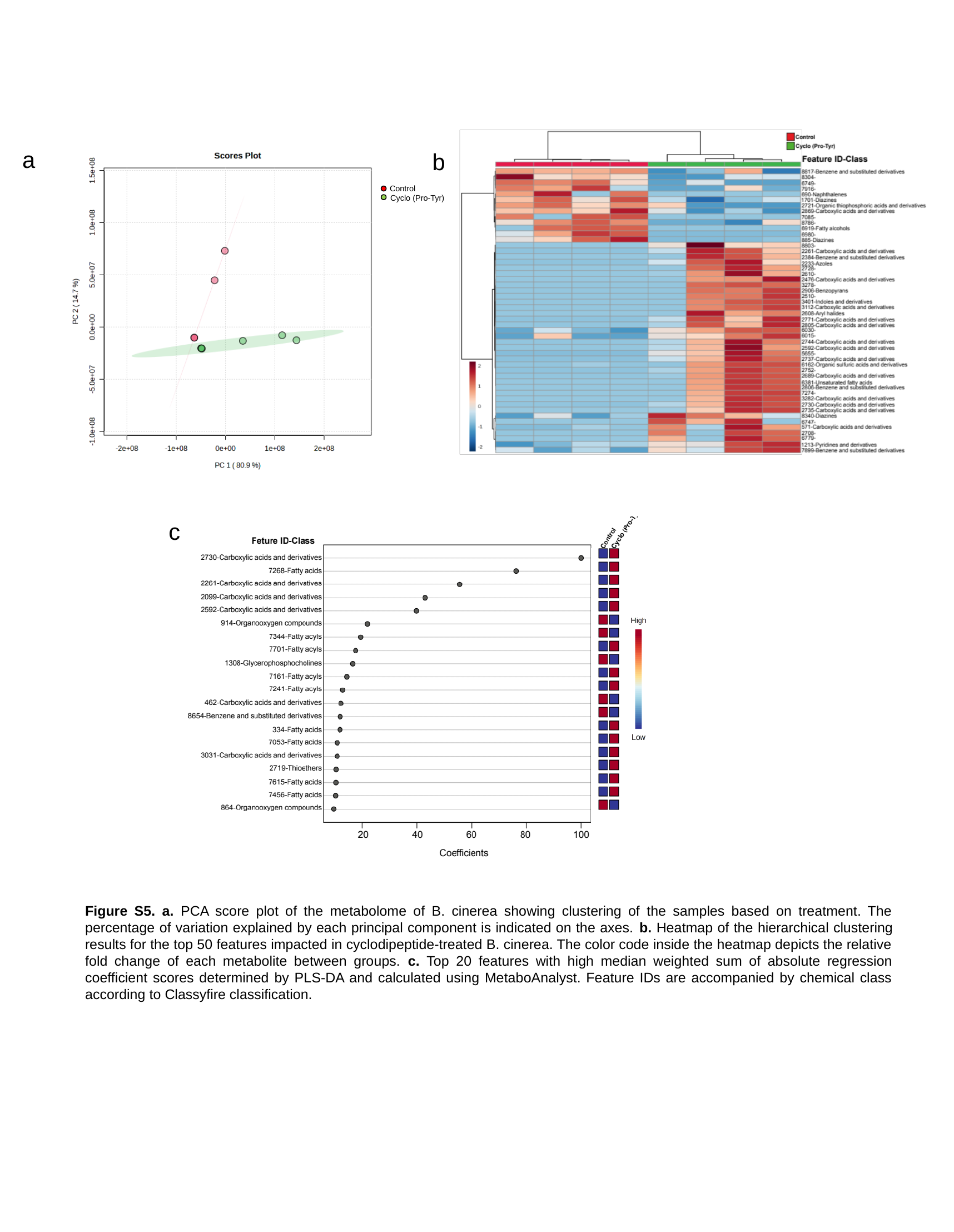

a
b
Control
Cyclo (Pro-Tyr)
c
Figure S5. a. PCA score plot of the metabolome of B. cinerea showing clustering of the samples based on treatment. The percentage of variation explained by each principal component is indicated on the axes. b. Heatmap of the hierarchical clustering results for the top 50 features impacted in cyclodipeptide-treated B. cinerea. The color code inside the heatmap depicts the relative fold change of each metabolite between groups. c. Top 20 features with high median weighted sum of absolute regression coefficient scores determined by PLS-DA and calculated using MetaboAnalyst. Feature IDs are accompanied by chemical class according to Classyfire classification.

### Slide 6
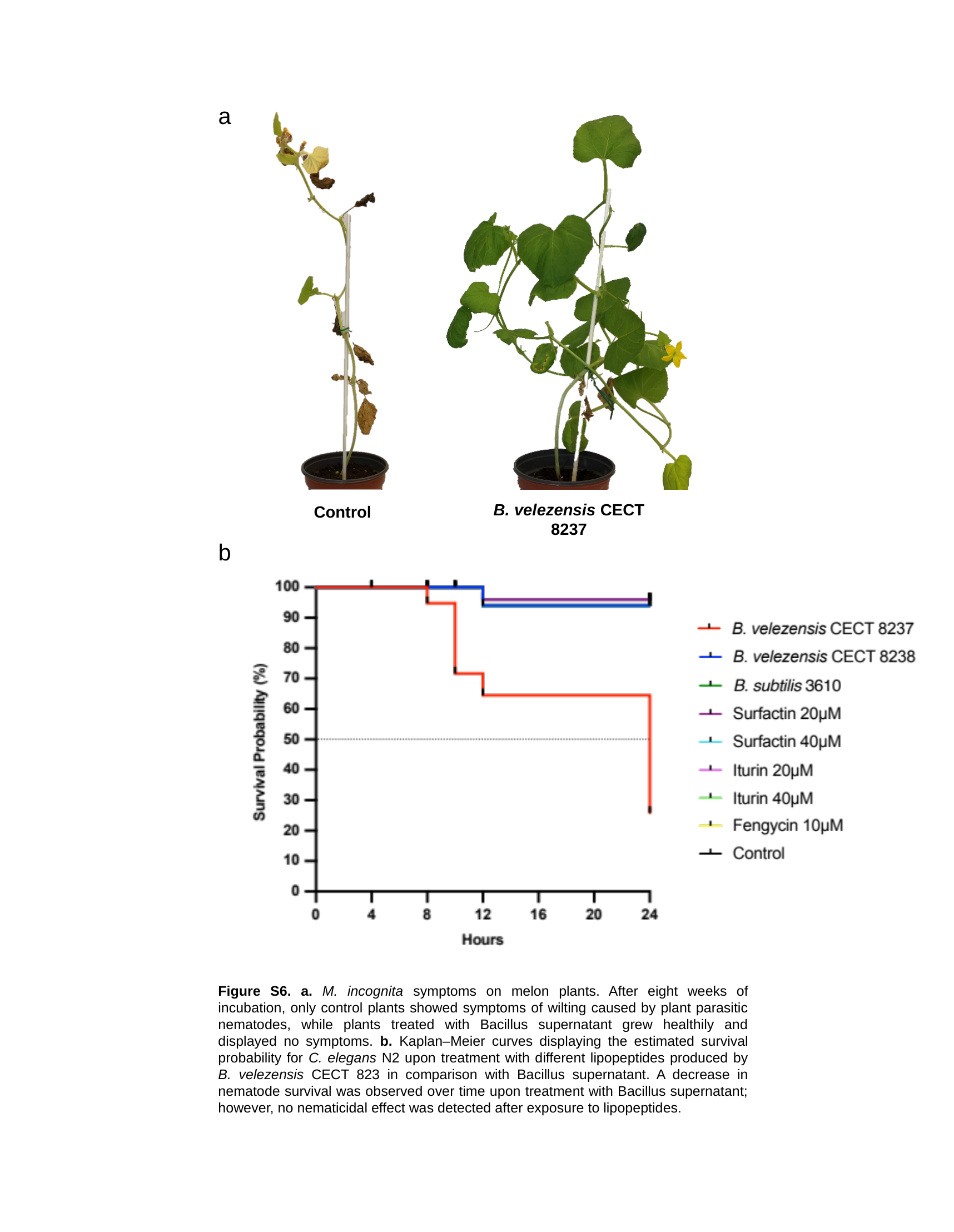

a
B. velezensis CECT 8237
Control
b
Figure S6. a. M. incognita symptoms on melon plants. After eight weeks of incubation, only control plants showed symptoms of wilting caused by plant parasitic nematodes, while plants treated with Bacillus supernatant grew healthily and displayed no symptoms. b. Kaplan‒Meier curves displaying the estimated survival probability for C. elegans N2 upon treatment with different lipopeptides produced by B. velezensis CECT 823 in comparison with Bacillus supernatant. A decrease in nematode survival was observed over time upon treatment with Bacillus supernatant; however, no nematicidal effect was detected after exposure to lipopeptides.

### Slide 7
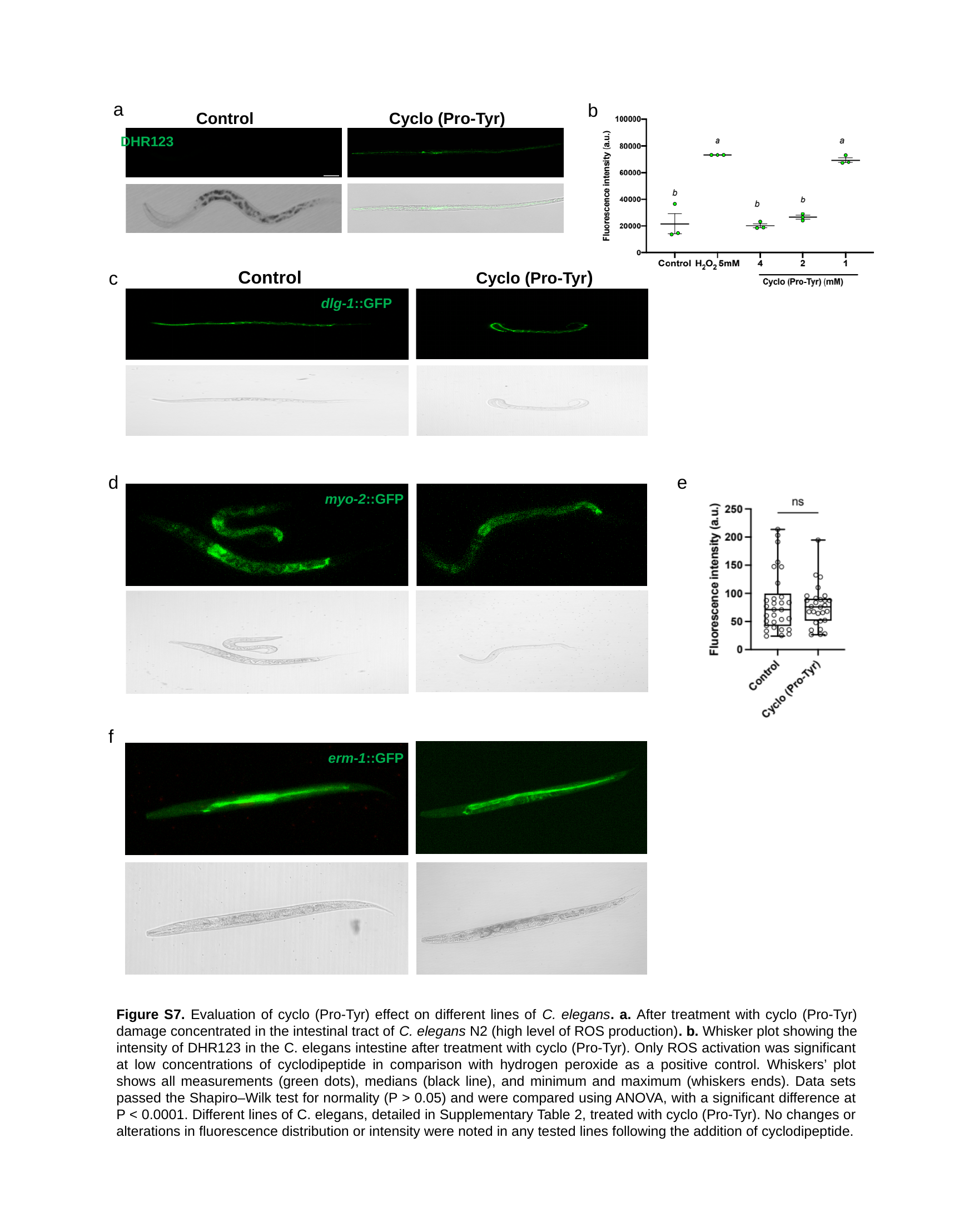

a
b
Control
Cyclo (Pro-Tyr)
DHR123
Control
Cyclo (Pro-Tyr)
dlg-1::GFP
c
d
e
myo-2::GFP
f
erm-1::GFP
Figure S7. Evaluation of cyclo (Pro-Tyr) effect on different lines of C. elegans. a. After treatment with cyclo (Pro-Tyr) damage concentrated in the intestinal tract of C. elegans N2 (high level of ROS production). b. Whisker plot showing the intensity of DHR123 in the C. elegans intestine after treatment with cyclo (Pro-Tyr). Only ROS activation was significant at low concentrations of cyclodipeptide in comparison with hydrogen peroxide as a positive control. Whiskers’ plot shows all measurements (green dots), medians (black line), and minimum and maximum (whiskers ends). Data sets passed the Shapiro‒Wilk test for normality (P > 0.05) and were compared using ANOVA, with a significant difference at P < 0.0001. Different lines of C. elegans, detailed in Supplementary Table 2, treated with cyclo (Pro-Tyr). No changes or alterations in fluorescence distribution or intensity were noted in any tested lines following the addition of cyclodipeptide.

### Slide 8
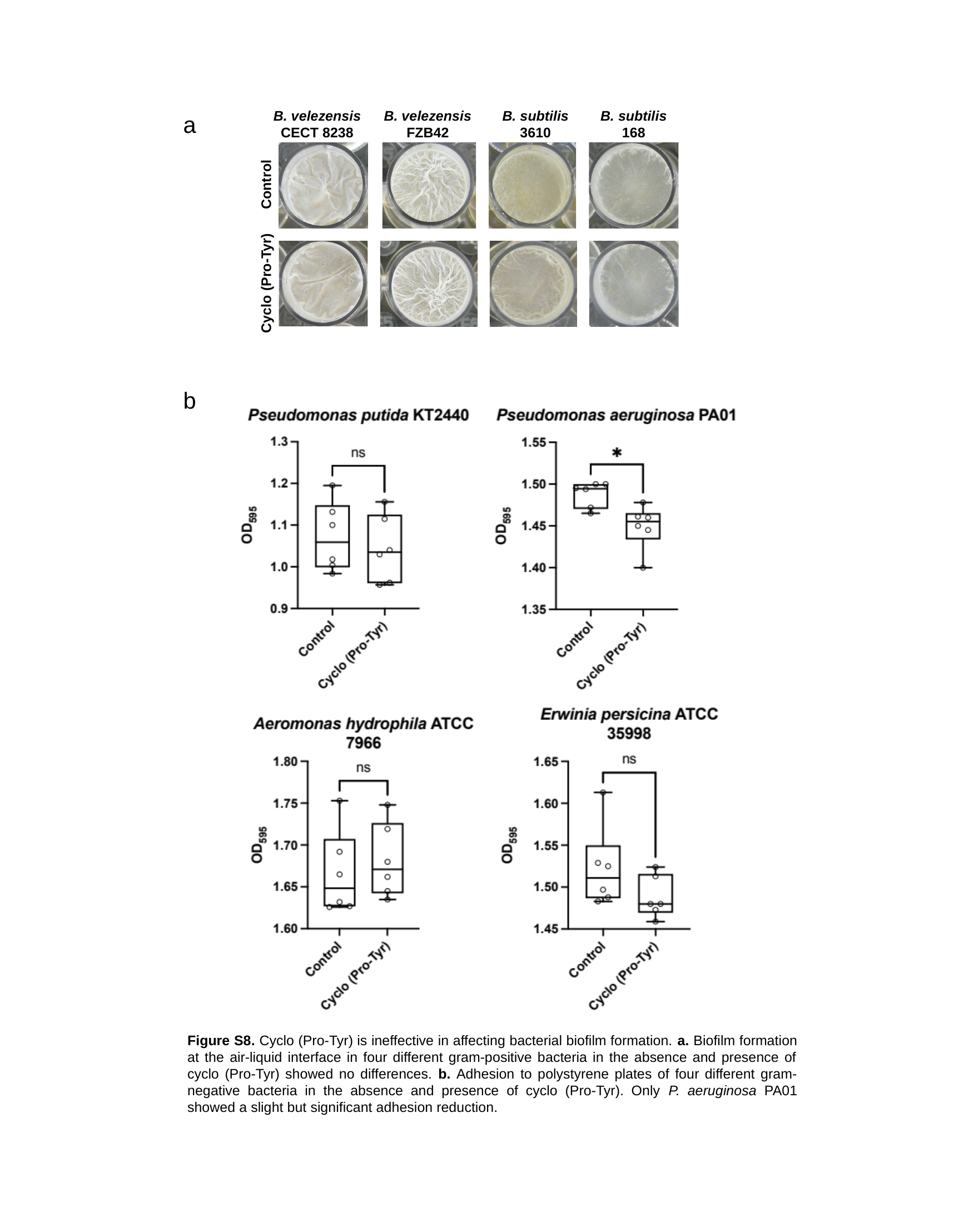

B. velezensis CECT 8238
B. velezensis FZB42
B. subtilis 3610
B. subtilis 168
Control
Cyclo (Pro-Tyr)
a
b
Figure S8. Cyclo (Pro-Tyr) is ineffective in affecting bacterial biofilm formation. a. Biofilm formation at the air-liquid interface in four different gram-positive bacteria in the absence and presence of cyclo (Pro-Tyr) showed no differences. b. Adhesion to polystyrene plates of four different gram-negative bacteria in the absence and presence of cyclo (Pro-Tyr). Only P. aeruginosa PA01 showed a slight but significant adhesion reduction.

### Slide 9
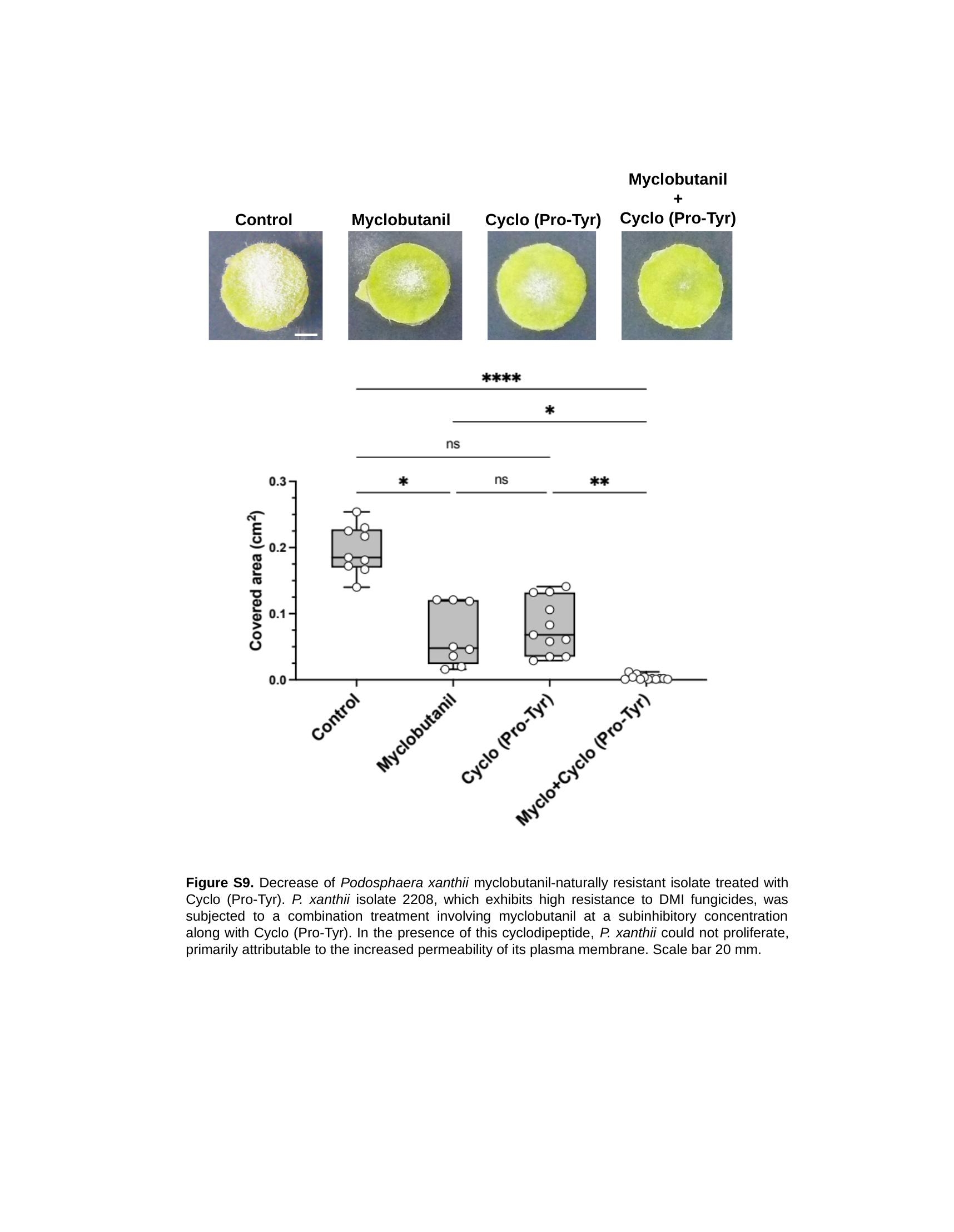

Myclobutanil
+
Cyclo (Pro-Tyr)
Control
Myclobutanil
Cyclo (Pro-Tyr)
Figure S9. Decrease of Podosphaera xanthii myclobutanil-naturally resistant isolate treated with Cyclo (Pro-Tyr). P. xanthii isolate 2208, which exhibits high resistance to DMI fungicides, was subjected to a combination treatment involving myclobutanil at a subinhibitory concentration along with Cyclo (Pro-Tyr). In the presence of this cyclodipeptide, P. xanthii could not proliferate, primarily attributable to the increased permeability of its plasma membrane. Scale bar 20 mm.

### Slide 10
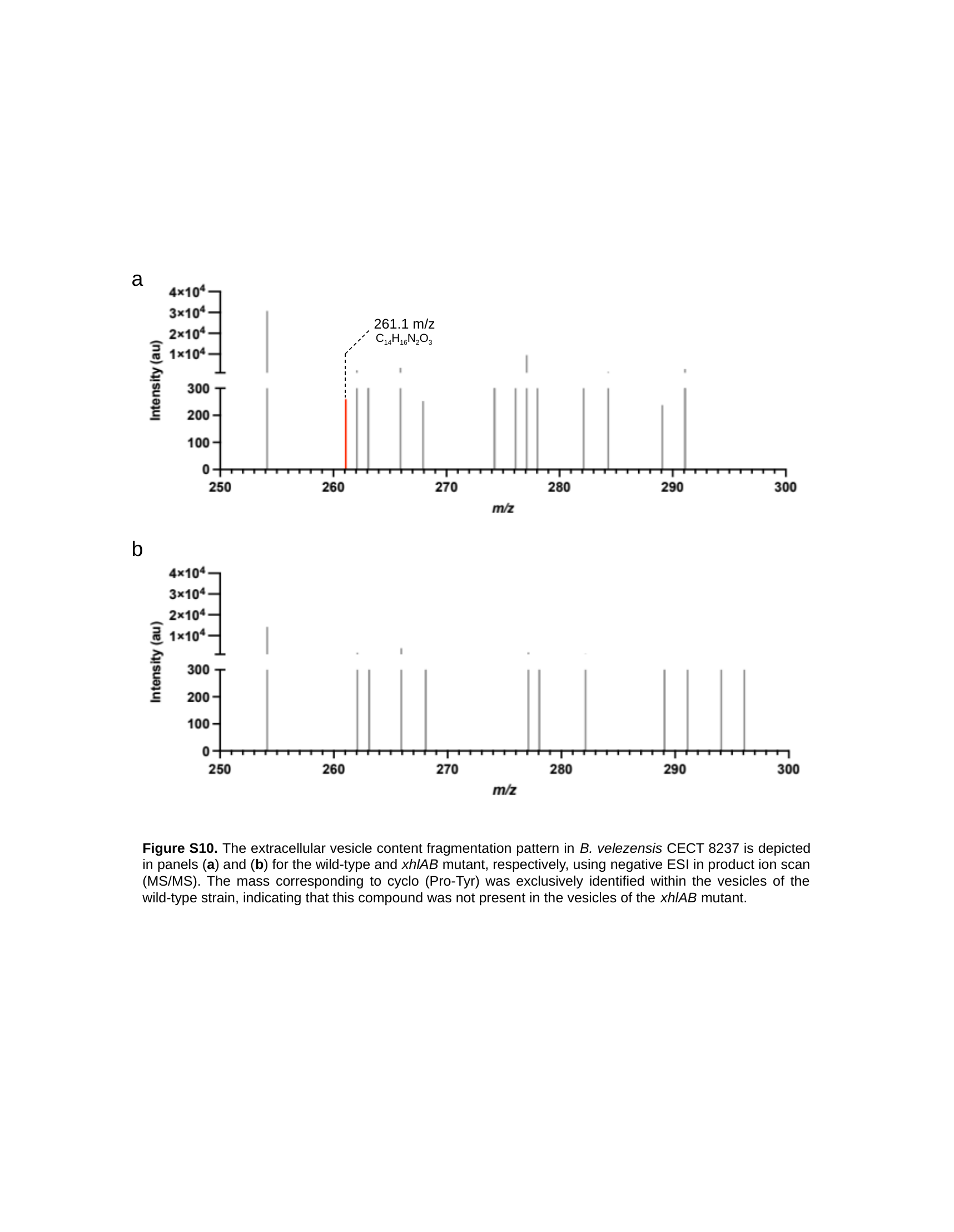

a
261.1 m/z
C_14_H_16_N_2_O_3_
b
Figure S10. The extracellular vesicle content fragmentation pattern in B. velezensis CECT 8237 is depicted in panels (a) and (b) for the wild-type and xhlAB mutant, respectively, using negative ESI in product ion scan (MS/MS). The mass corresponding to cyclo (Pro-Tyr) was exclusively identified within the vesicles of the wild-type strain, indicating that this compound was not present in the vesicles of the xhlAB mutant.

### Slide 11
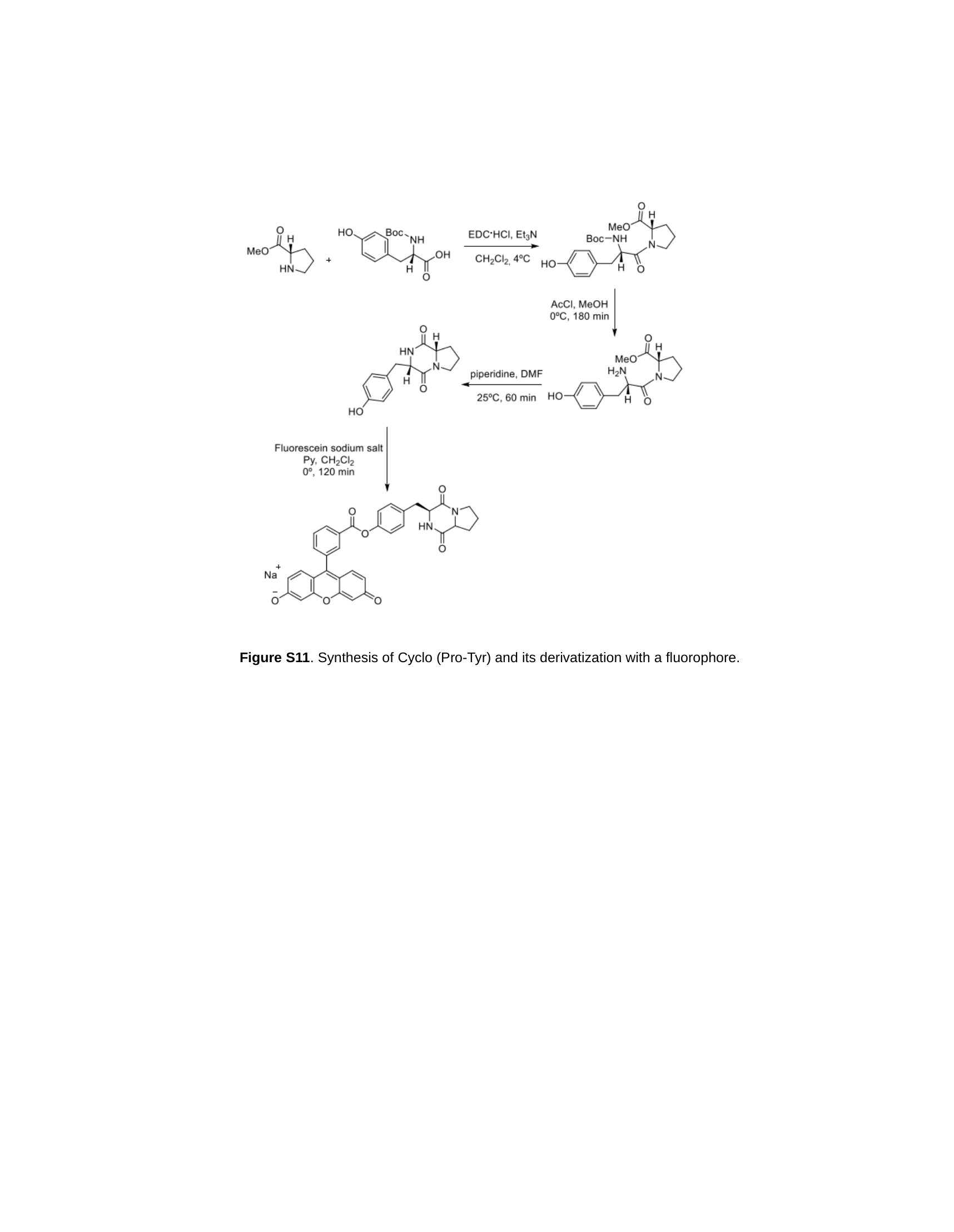

Figure S11. Synthesis of Cyclo (Pro-Tyr) and its derivatization with a fluorophore.

### Slide 12
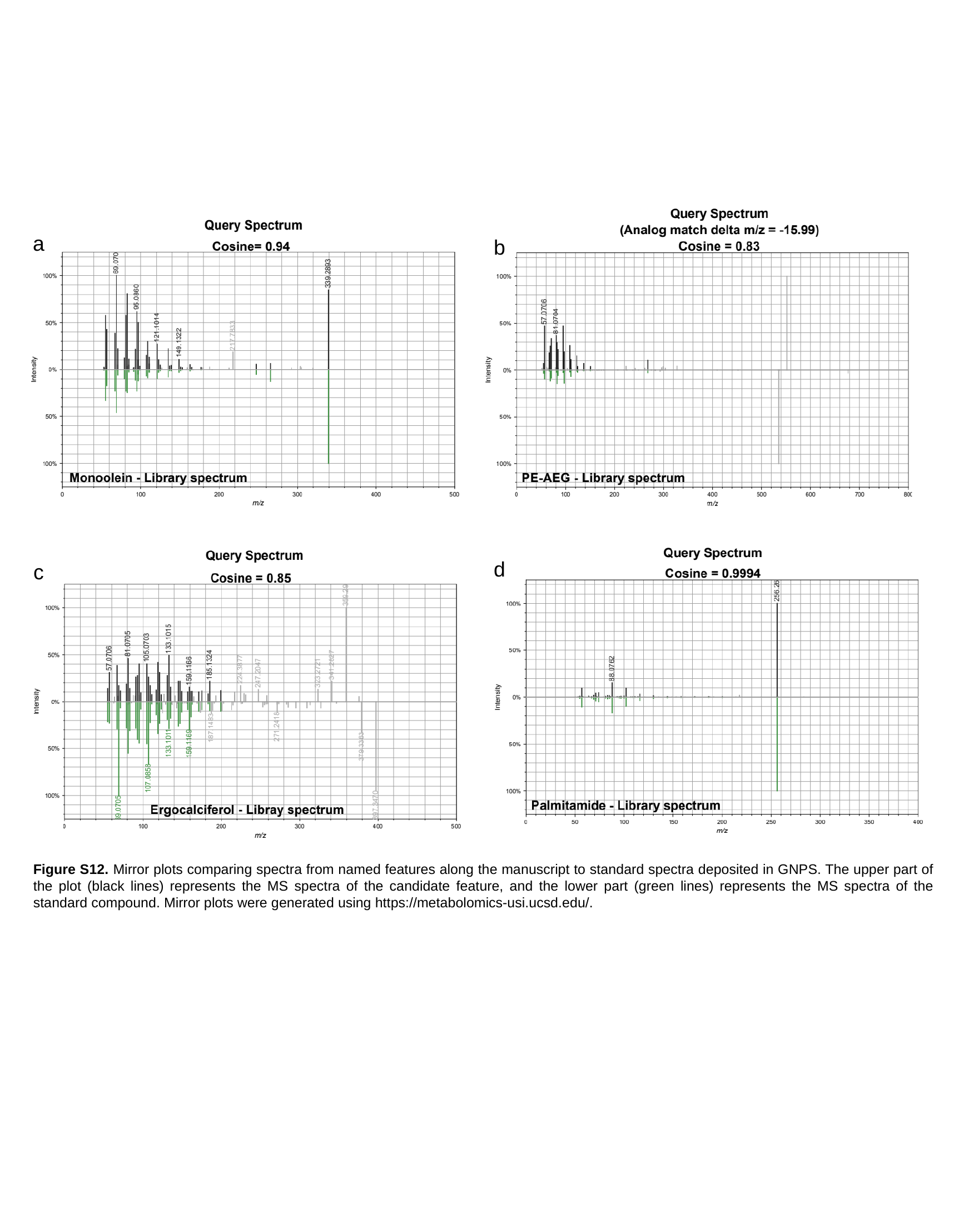

a
b
d
c
Figure S12. Mirror plots comparing spectra from named features along the manuscript to standard spectra deposited in GNPS. The upper part of the plot (black lines) represents the MS spectra of the candidate feature, and the lower part (green lines) represents the MS spectra of the standard compound. Mirror plots were generated using https://metabolomics-usi.ucsd.edu/.
